# Neuroanatomy of substantia nigra and ventral tegmental area dopaminergic, and dorsal raphe serotonergic circuits in the human brain using T1-weighted and diffusion magnetic resonance imaging: A morphometric pilot study with estimate of reliability

**DOI:** 10.64898/2026.02.12.705574

**Authors:** Poliana Hartung Toppa, Richard J. Rushmore, Kayley Haggerty, George Papadimitriou, Darin Dougherty, Marek Kubicki, José Luis González-Mora, Stefano Pallanti, Agustin Castañeyra-Perdomo, Edward Yeterian, Nikos Makris

**Affiliations:** Center for Morphometric Analysis, Departments of Psychiatry and Neurology, Athinoula A. Martinos Center for Biomedical Imaging, Massachusetts General Hospital, Harvard Medical School, Boston, Massachusetts 02129, United States; Department of Anatomy and Neurobiology, Boston University School of Medicine, Boston, Massachusetts 02118, United States; Psychiatry Neuroimaging Laboratory, Department of Psychiatry, Brigham and Women’s Hospital, Harvard Medical School, Boston, Massachusetts 02115, United States; Department of Psychiatry, Massachusetts General Hospital, Harvard Medical School, Boston, MA, United States; Universidad de La Laguna, Área de Anatomía y Fisiología, Departamento de Ciencias Médicas Básicas. Facultad de Ciencias de la Salud, San Cristobal de La Laguna 38000, Tenerife, Spain; Universidad de La Laguna, Instituto Universitario de Neurosciencias, Facultad de Ciencias de la Salud, San Cristobal de La Laguna 38000, Tenerife, Spain; Department of Psychiatry and Behavioral Science, Albert Einstein College of Medicine, Bronx, New York, United States; Institute for Neuroscience, Florence, Italy; Department of Psychology, Colby College, Waterville, Maine 04901, United States

**Author notes:** The authors contributed equally to this paper (Poliana Hartung Toppa, Richard J. Rushmore). The authors contributed equally to this paper (Edward Yeterian, Nikos Makris).

**Keywords:** diffusion MRI, structural connectivity, dopamine, serotonin, major depressive disorder, COVID-19, long COVID, dorsolateral prefrontal cortex, cerebellum

## Abstract

**Introduction:** We present here a methodology for morphometric analysis of the substantia nigra (SN), the ventral tegmental area (VTA), the dorsal raphe nucleus (DRN) and their respective structural brain circuits.

**Methods:** Our analyses were based on multimodal T1-weighted MRI and diffusion MRI (dMRI) segmentation and tractography in 12 human subjects drawn from the Human Connectome Project (HCP) repository.

**Results:** We were able to demonstrate strong connections of the SN, VTA and DRN with several brain regions, in particular the dorsolateral prefrontal cortex (DLPFC) and the cerebellum. More specifically, we created comprehensive visualizations of the SN and VTA dopaminergic as well as the DRN serotonergic structural circuits in the human brain, which, although preliminary, demonstrate the potential of multimodal neuroimaging to investigate these circuits quantitatively in clinical conditions. Finally, we created a pilot dataset for the most frequently observed structural connections, specifically those that were present more than 92% of the time among all subjects. <u>Discussion</u> This pilot morphometric report examines the structural circuits of the SN, VTA and DRN, which are critically involved in several biobehaviors and clinical conditions such as addiction, stress, Parkinson’s disease (PD), schizophrenia, obsessive-compulsive disorder, post-traumatic stress disorder, attention deficit hyperactivity disorder, mood disorders, COVID-19 and long COVID. Importantly, the strong structural connectivity of the DLPFC and cerebellum with the SN, VTA and DRN is expected to be a potential target of noninvasive neuromodulation treatments in neuropsychiatry. Our findings demonstrate the potential of current clinical multimodal neuroimaging to delineate the dopaminergic (DA) and serotonergic (5-HT) circuits in the human brain in clinical conditions.

## Introduction

Monoamines are a group of neurotransmitters in the brain, which include catecholamines such as dopamine (DA) and indolamines such as serotonin or 5-hydroxytryptamine (5-HT). Catecholamines (or central catecholamines) include norepinephrine (NE) (or noradrenaline) and epinephrine (E) (or adrenaline) as well as dopamine. The topographical and structural organization of the monoamine-containing neurons shows a notable consistency in the mammalian brain. These cell groups are located principally within the brainstem and the hypothalamus. In particular, DA is produced in the midbrain and the hypothalamus, whereas 5-HT is produced in the mesencephalon (midbrain), pons and medulla (Nieuwenhuys, 1985; Cooper et al., 2003; Nolte, 2009). The precise topography of monoamine circuits in the human brain is largely unknown and is derived primarily from animal experimental studies that began in the 1960s (Nieuwenhuys, 1985). Using current multimodal neuroimaging our group has initiated a systematic program of research on monoamine structural pathways in postmortem and in vivo human brains (Rushmore et al., 2020; Castañeyra-Perdomo et al., 2024; Kikinis et al., 2024; Makris et al., 2024; Toppa et al., 2025). To date, we have delineated the noradrenergic pathways as reported in Makris et al. (2024) and Toppa et al. (2025). These studies have been aimed at integrating chemical neuroanatomy with traditional neuroanatomy in the human brain, as pioneered and elaborated upon by Nieuwenhuys (1985) and further developed by our group (e.g., (Rushmore et al., 2020; Makris et al., 2024). The focus of the present report is on the structural connectivity of dopaminergic and serotonergic brain circuitry, to complement and advance our prior studies of the neurochemical anatomy of the human brain (Rushmore et al., 2020; Castañeyra-Perdomo et al., 2024; Kikinis et al., 2024; Makris et al., 2024; Toppa et al., 2025).

The biobehavioral and clinical effects of dopamine are central in reward function, motivation and mood regulation. Moreover, DA plays a role in cognitive processing including memory and attention as well as many other bodily functions, especially movement. Thus, the understanding of dopaminergic human brain circuits is key in basic and clinical neuroscience (Stevens, 1973; Bird et al., 1979; Stevens et al., 1979; Reynolds, 1983; Koob, 1992, 1999; Blum et al., 1996; Breiter et al., 1997; Bowirrat and Oscar-Berman, 2005; Berridge, 2007; Koob and Volkow, 2010). Likewise, 5-HT is associated with key brain and bodily functions, such as mood, sleep, digestion, blood coagulation and wound healing. Unbalance of serotonin in the brain and body underlies clinical conditions such as depression and anxiety. Both DA and 5-HT play a role in sexual desire (Taber et al., 1960; Dahlström and Fuxe, 1964; Braak, 1970; Sladek and Walker, 1977; Wiklund and Björklund, 1980; Takeuchi et al., 1982; Bowker et al., 1983; Felten and Sladek, 1983). The dopaminergic and serotonergic systems have specific cells of origin in the brainstem, which give rise to axonal fiber pathways that course via specific topographic trajectories toward their terminations throughout the brain (Nieuwenhuys, 1985; Cooper et al., 2003). Furthermore, given the nature of clinical practice, which necessitates both molecular-metabolic and anatomical-histological knowledge and clarity associated with therapeutic interventions, the identification of these neurochemical systems and the ability to monitor their structure and function in health and disease is a matter of great clinical relevance. To this end, exact knowledge of dopaminergic and serotonergic brain circuits is of particular importance for targeting precision care therapeutic interventions and monitoring their effectiveness. Importantly, the advent of neuroimaging has been of great advantage, given its non-invasive, in vivo nature that has resulted in the development of novel research and clinical opportunities. In principle, non-invasive transcranial magnetic stimulation (TMS) and invasive deep brain stimulation (DBS) therapeutic approaches in clinical neuroscience can become much more precise and tailored to individual brains using modern neuroimaging approaches. Detailed topographic mapping of neurochemical circuits is critical to ensuring precise and specific targeting using TMS and DBS. It is in this context that the present study of DA and 5-HT structural circuitry in the human brain aims to delineate precisely these potential target circuits and thus contribute toward making these therapies more accurate and effective. We have elaborated on this important topic in a recent publication by Lioumis et al. (Lioumis et al., 2025).

Given the complex anatomy of the brainstem and the hypothalamus as well as the spatial limitations these brain regions present for neuroimaging, clinical structural neuroimaging investigations of fine-grained brainstem and hypothalamic anatomy are extremely challenging (Calabrese et al., 2015). In this regard, multimodal neuroimaging allows the identification of brain gray matter structures as well as the fiber tracts that interconnect them. It should be noted that we use the term “multimodal” imaging to refer to the use of two or more imaging modalities whose unique properties complement each other and are necessary and sufficient for the task to be addressed. Specifically, in the present study T1-weighted magnetic resonance imaging (MRI) morphometry enables the segmentation of brainstem, diencephalic and other cerebral central gray matter and cortical regions of interest (ROIs), while diffusion MRI (dMRI) tractography permits the delineation of white matter fiber pathways. Thus, the brain circuits associated with the DA and 5-HT systems, which are constituted by gray matter cortical regions, subcortical nuclei and their associated axonal white matter fibers, can be sampled and measured reliably using current neuroimaging.

In the present MRI-based neuroanatomy study we report results on the substantia nigra (SN) and ventral tegmental area (VTA) dopaminergic circuits and the dorsal raphe nucleus (DRN) serotonergic circuits in 12 human subjects comprising a balanced cohort of six females and six males matched in age drawn from the Human Connectome Project (HCP) repository (Van Essen et al., 2013). To this end, we used a novel methodology that combined structural T1-weighted MRI morphometric and dMRI tractographic analyses. Specifically, we reconstructed the mesostriatal, mesolimbic and mesocortical DA projection systems in three-dimensional space. We also delineated the connections of the dorsal raphe nucleus, which is the richest source of serotonergic connections in the human brain. More specifically, we investigated the connections of the SN, VTA and DRN guided by animal experimental literature following the Pandya comparative extrapolation principle (Makris et al., 2023b) as well as by the more limited human literature in this area (e.g., (Edlow and Wu, 2012; Edlow et al., 2016, 2021; García-Gomar et al., 2022b; Skandalakis et al., 2024). We explored structural connections of the SN, VTA and DRN with the prefrontal cortex and cingulate gyrus in view of the considerable expansion of these regions in the human brain and their critical relevance in executive function and cognitive control. In addition, we measured biophysical parameters of average fractional anisotropy (FA) (Basser, 2004), axial diffusivity (AD) (Song et al., 2003) and radial diffusivity (RD) (Song et al., 2002, 2003) of all fiber tracts in the 12 healthy human subjects. We produced visualizations of the SN and VTA dopaminergic as well as the DRN serotonergic structural circuits in the human brain. We also reported biophysical parameters for the most frequently observed structural connections, specifically those that were present in 92% or more of all subjects. We expect that the study of brain circuits addressing both their neurochemical architecture and their precise location and connectivity in terms of specific origins and terminations will enhance our understanding of structural-functional relationships, a matter of great value in elucidating brain organization.

## Materials and Methods

### Methodology

The methodology used in the present study was performed in two consecutive steps as illustrated in detail in **Figure 1**. In the <u>first step,</u> we defined and segmented the SN, VTA, and the DRN ROIs as follows. 1) We segmented the SN ROI using the b0 series of the dMRI acquisition. The SN is located in the diencephalon and the midbrain (e.g., (Carleton and Carpenter, 1984; Nieuwenhuys et al., 2007). Using the b0 spherical mean (Legarreta et al., 2025) we were able to directly visualize and identify the SN. Specifically, diffusion sequences were available at a 1.25mm^3^ isotropic resolution at b value shells of 1000, 2000, and 3000s/mm^2^ acquired at 270 directions. The spherical mean for each shell across all gradient encoding directions was calculated, then merged to produce the contrast for which the region corresponding to the SN exhibited a specific decrease in intensity relative to surrounding structures. In the present MRI analysis, we were unable to subdivide the SN into pars compacta and pars reticulata, given that b0 dMRI and T1-weighted MRI images do not allow distinguishing between the two parts of the SN with clarity. Thus, we sampled the SN as a single ROI, which was clearly distinguishable from neighboring structures such as the red nucleus (RN) and the cerebral peduncle (CP) (**Fig. 1Ai-ii**). We outlined the SN using the “threshold” tool of 3D Slicer’s neurosegmentation module, which allows fine-tuning of the borders of the structure (**Fig. 1Aiii**). This module allows the implementation of the histogram method (Worth et al., 1998) to determine the thresholds needed for the segmentation of the structural ROIs (**Fig. 1Aiii**). This is performed by sampling representative voxels around the border of the SN, as indicated by the circle in **Figure 1Aiii**. The histogram method is critically important for the reliable segmentation of anatomical ROIs in morphometric analysis (Filipek et al., 1994; Worth et al., 1998). This methodological issue has been addressed in detail since the inception of morphometric analysis by our group (e.g., (Rademacher et al., 1992; Filipek et al., 1994; Rushmore et al., 2022). This procedure used the threshold tool as shown in **Figure 1Aiv**. This was done on a complete series of axial sections from the superiormost to the inferiormost sections on which the SN was identified. We then produced 3D reconstructions of the SN ROI and ensured that they were consistent with the known neuroanatomical location of the SN (**Fig. 1Av**). 2) We segmented the VTA ROI using T1-weighted MRI axial images as shown in **Figure 1Bi**. It should be noted that the outlines of the SN ROI are a prerequisite for the VTA ROI segmentation given that the SN determines the lateral and anterior borders of the VTA. Thus, the SN outline was transferred to the corresponding T1-weighted images from the co-registered b0 images (e.g., **Fig. 1Bi-ii**) on which the original SN ROI segmentation was performed. The VTA ROI is located in the peduncular tegmentum and, more precisely, lies adjacent to the midline borders of the interpeduncular fossa (ipf), maintaining a medial position with respect to the SN (**Fig. 1Bi-ii**). Its medial border is the interpeduncular fossa (**Fig. 1Bi-ii**), whereas its lateral border approximates the medial aspect of the SN (**Fig. 1Bii**). Anatomical knowledge of the VTA’s topographic relationships with the SN is critical for the definition of the VTA ROI given that the VTA is not detectable by current clinical T1-weighted MRI or dMRI protocols. Specifically, the VTA ROI is located in the area enclosed by the SN laterally and anteriorly, and medially by the midbrain exterior margin at the ipf. Its superior border is defined by the axial plane at the inferiormost point of the inferior colliculus as shown by the dotted white horizontal line in **Figure 1Biii**. Finally, its inferior border is the axial plane at the inferiormost point of the interpeduncular fossa at the pons-midbrain transition as shown by the solid white horizontal line in **Figure 1Biii**. 3) The DRN ROI was based on our brainstem parcellation methodology as described by Makris et al. (Makris et al., 2024). Briefly, the pons was divided into upper (P1) and lower (P2) parts by an axial plane parallel to the ACPC (bicommissural) line passing through the mid-pontine point of the basilar pons. The superior border of P1 is defined by a plane that extends from the inferiormost point of the interpeduncular fossa, which, ventrally, is the junction between the pons and the midbrain, to the caudal endpoint of the inferior colliculus. The P1 and P2 parts were further subdivided into an anterior (P1a, P2a) and a posterior (P1p, P2p) parcel by a transverse line between the basilar-tegmental notch on the lateral aspects of the pons. The DRN ROI segmentation was performed in a series of T1-weighted MRI axial sections within the posterior upper part of the pons and the posterior part of the lower midbrain as shown in **Figure 1Ci-iv**. Its medial border was the midline, whereas its lateral border was, by convention, a vertical line drawn laterally at a distance 1/8^th^ of the width of P1p from the midline (which in these data corresponded to three 0.7mm isotropic voxels from the midline) as shown in **Figure 1Ciii, iv**. The anterior border of the DRN ROI was set at half the distance from the border between P1a and P1p in the pons (DaSilva et al., 2002; Makris et al., 2024; Toppa et al., 2025) and M2 in the midbrain (DaSilva et al., 2002; Makris et al., 2024; Toppa et al., 2025) (**Figure 1Ci-iv**). The inferior border was the line joining the inferiormost point of the interpeduncular fossa with the transition point of the aqueduct with the 4^th^ ventricle (**Figure 1Ci, ii**). Finally, its superior border was defined by a line joining the mid-collicular point, i.e., the junction of the superior and inferior colliculi, with the midpoint of a line defining the border between the pons and the midbrain (i.e., the line between the interpeduncular fossa and the inferiormost point of the cerebral aqueduct (**Figure 1Cii)**. Finally, representative 3D reconstructions of the left, right and bilateral SN, VTA and DRN ROIs were generated, as shown in **Figures 1Av, 1Biv-vi and 1Cv-vii**, respectively.

**Figure 1.**
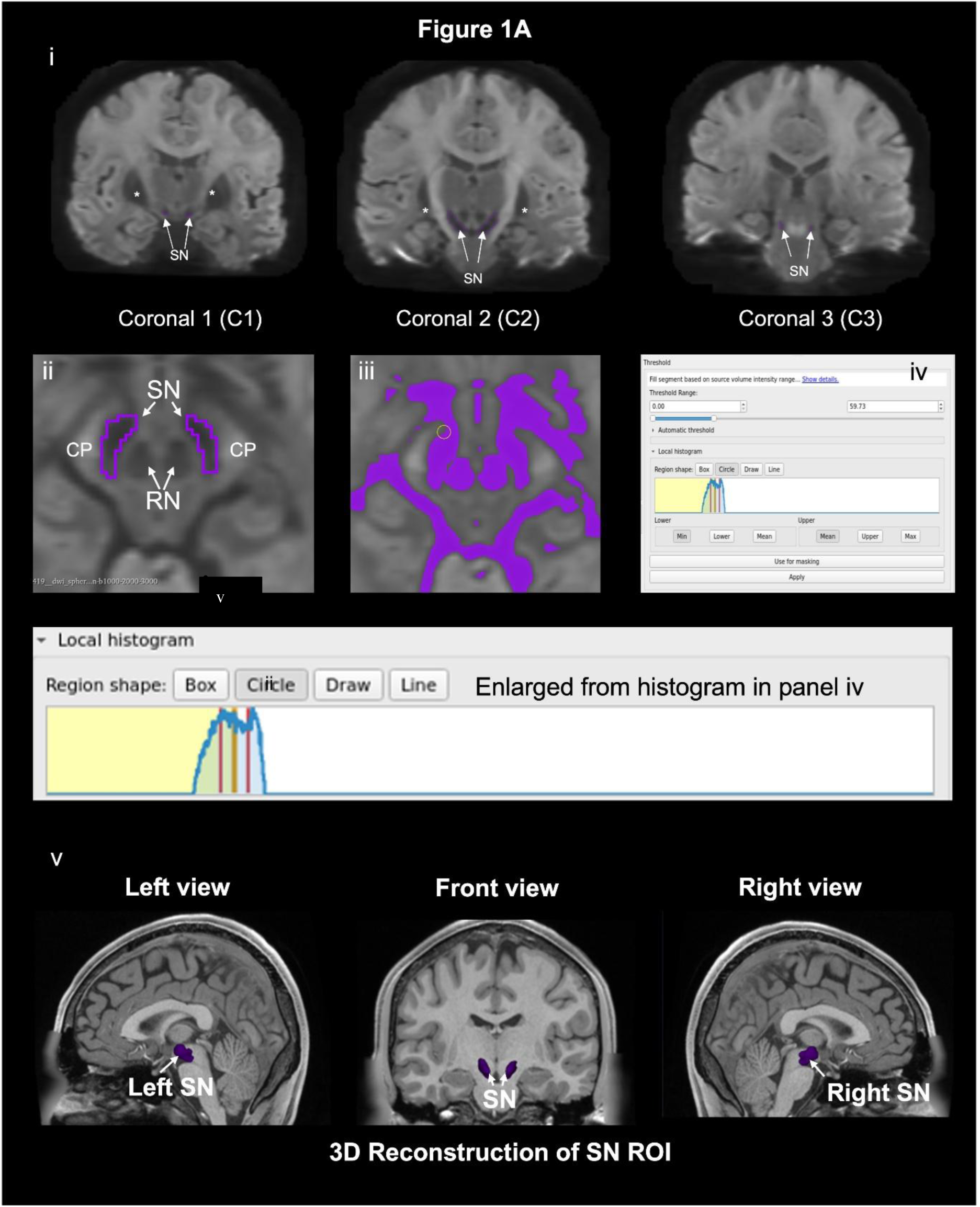
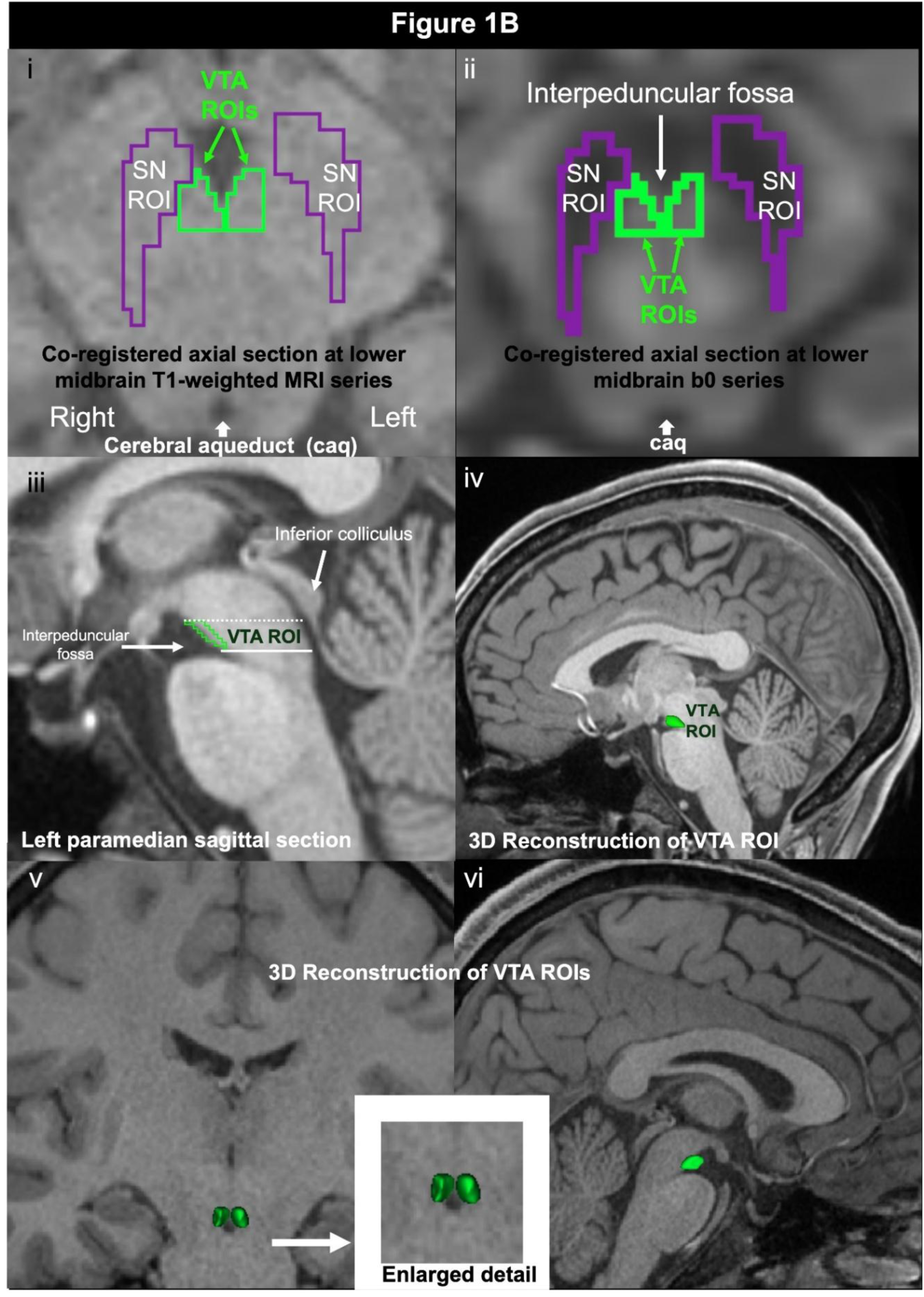
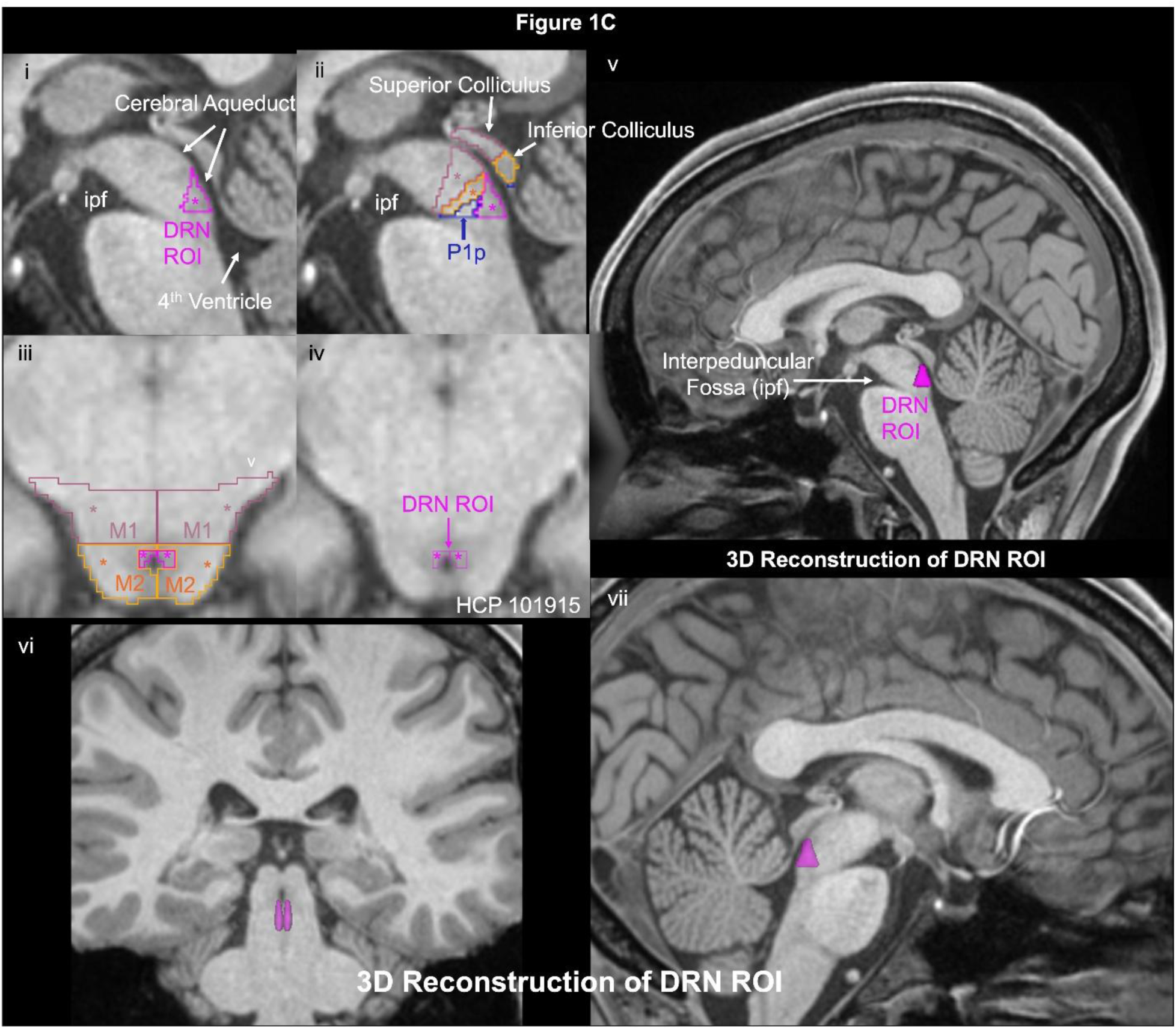
Segmentation methodology of the substantia nigra (SN), ventral tegmental area (VTA) and dorsal raphe nucleus (DRN) regions of interest (ROIs). Please see detailed methodology in the methods section. In **Figure 1A**, the SN was sampled as a single ROI, which was clearly distinguishable from neighboring structures such as the red nucleus (RN) and the cerebral peduncle (CP) using the b0 series of the dMRI acquisition (**Fig. 1Ai-ii**). To this end, the “threshold” tool of the neurosegmentation module of 3D Slicer was used (**Fig. 1Aiii**). This module allows the implementation of the histogram method to determine the thresholds needed for the segmentation of the structural ROIs (**Fig. 1Aiii-iv**). This procedure used the threshold tool as shown in **Figure 1Aiv**. This was done on a complete series of axial sections from the superiormost to the inferiormost sections on which the SN was identified. We then produced 3D reconstructions of the SN ROI and ensured that they were consistent with the known neuroanatomical location of the SN (**Fig. 1Av**). In **Figure 1B**, the VTA ROI was segmented using T1-weighted MRI axial images as shown in **Figure 1Bi**. The VTA ROI lies adjacent to the midline borders of the interpeduncular fossa, maintaining a medial position with respect to the SN (**Fig. 1Bi-ii**). Its borders are elaborated upon in detail in the methods section as shown in **Figure 1Bi-iii**. As explained in the methods section, the outlines of the SN ROI are a prerequisite for the VTA ROI segmentation and thus were transferred to the corresponding T1-weighted images from the co-registered b0 images (e.g., **Fig. 1Bi-ii**) on which the original SN ROI segmentation was performed. In **Figure 1C** the DRN ROI segmentation is depicted in T1-weighted MRI axial sections within the posterior upper part of the pons and the posterior part of the lower midbrain (**Fig. 1Ci-iv**). The details of the segmentation are explained in the methods section. Representative 3D reconstructions of the left, right and bilateral SN, VTA and DRN ROIs were generated, as shown in **Figures 1Av, 1Biv-vi and 1Cv-vii**, respectively.

It should be noted that the segmentation method for the human brainstem has been reported and applied in several studies by our group (DaSilva et al., 2002; Rivas-Grajales et al., 2018; Kikinis et al., 2024). Segmentation was performed using the “neurosegmentation module” of the publicly available 3D Slicer software platform (www.slicer.org; (Rushmore et al., 2022). All cortical and subcortical ROIs in the cerebrum were derived using FreeSurfer (Fischl et al., 2002, 2004) as provided by the publicly available HCP website (https://www.humanconnectome.org) (Van Essen et al., 2013).

The SN, VTA and DRN ROIs were sampled and used as “seeds” for tractography in the second step of the analysis. The <u>second step</u> involved dMRI-based tractography for the delineation of the SN, VTA and DRN structural connectivity. The full list of connections investigated herein is presented in **Table 1**. These connections have been described extensively in the experimental animal literature and are detailed in the discussion section. It should be noted that given the paucity of human traditional anatomical studies addressing these structural connections, we followed the paradigm of the Pandya comparative extrapolation principle (Makris et al., 2023b) and the approach of MRI-based brain volumetrics (Caviness et al., 1999; Makris et al., 2023a) as we have adopted in several of our studies. Given the limited literature regarding contralateral structural connectivity of the SN, VTA and DRN in experimental animals and humans, we limited our study to their ipsilateral structural connectomes (e.g., (Nieuwenhuys, 1985)). White Matter Query Language (WMQL) was employed to automatically extract white matter fascicles from whole-brain tractography based on defined anatomical regions (Wassermann et al., 2013, 2016). Specifically, WMQL allowed the reconstruction of the virtual fiber tracts connecting each of the three “seed” ROIs with other brain regions. Subsequently, the Slicer dMRI module of the 3D Slicer software platform (www.slicer.org) was used for the visualization of the SN, VTA and DRN structural circuits. Finally, we performed minor manual editing using the Slicer dMRI module of the 3D Slicer software platform (www.slicer.org).

**Table 1:**
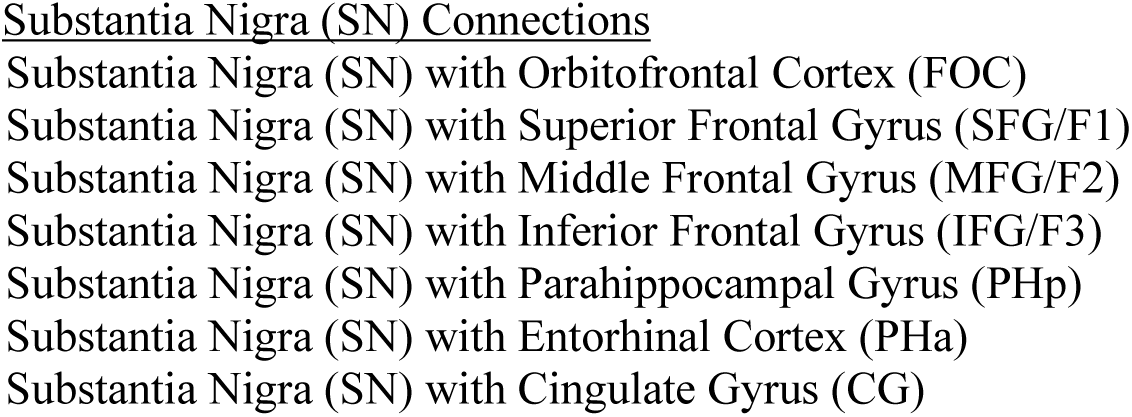
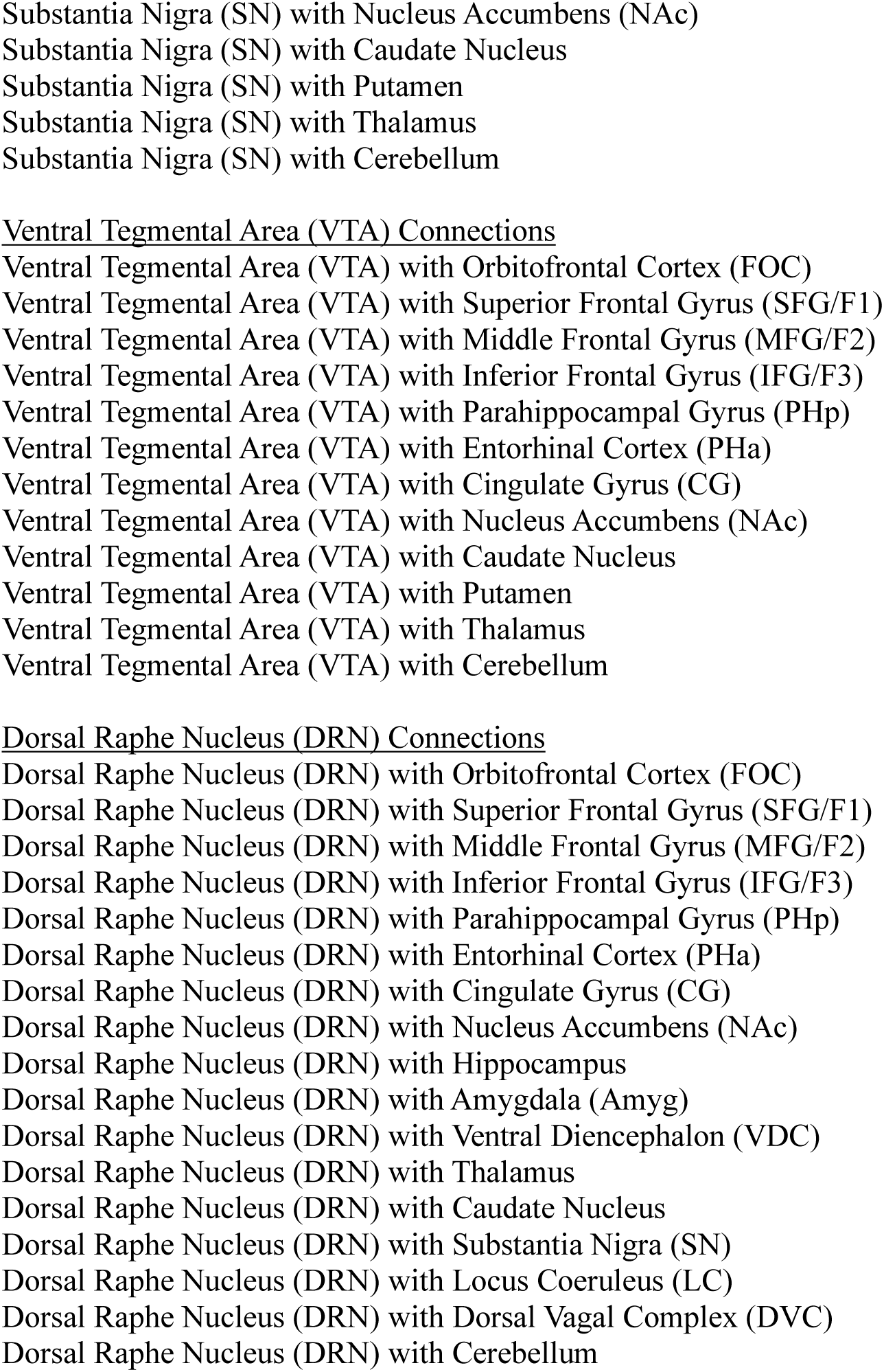
Structural connections delineated in this study.

### Reliability assessment

Before proceeding to the second step (described above), which included the tractographic reconstruction of fiber tracts, we first performed morphometric segmentation blindly in five representative subjects of our sample population. This was done by two well-trained image analysts who had background training in neuroanatomy. We computed intra-rater and inter-rater reliability measurements for the SN, VTA and DRN segmentations, using the Dice coefficient.

### Subjects

The 12 datasets analyzed herein were part of the publicly available HCP repository (https://www.humanconnectome.org) (Van Essen et al., 2013) and are listed in **Table 2**. All subjects were classified as healthy controls. The age range of the six females was 22 to 35 with an average age of 29.5 years. The age range of the six males was 22 to 35 with an average age of 28.9 years.

**Table 2:**
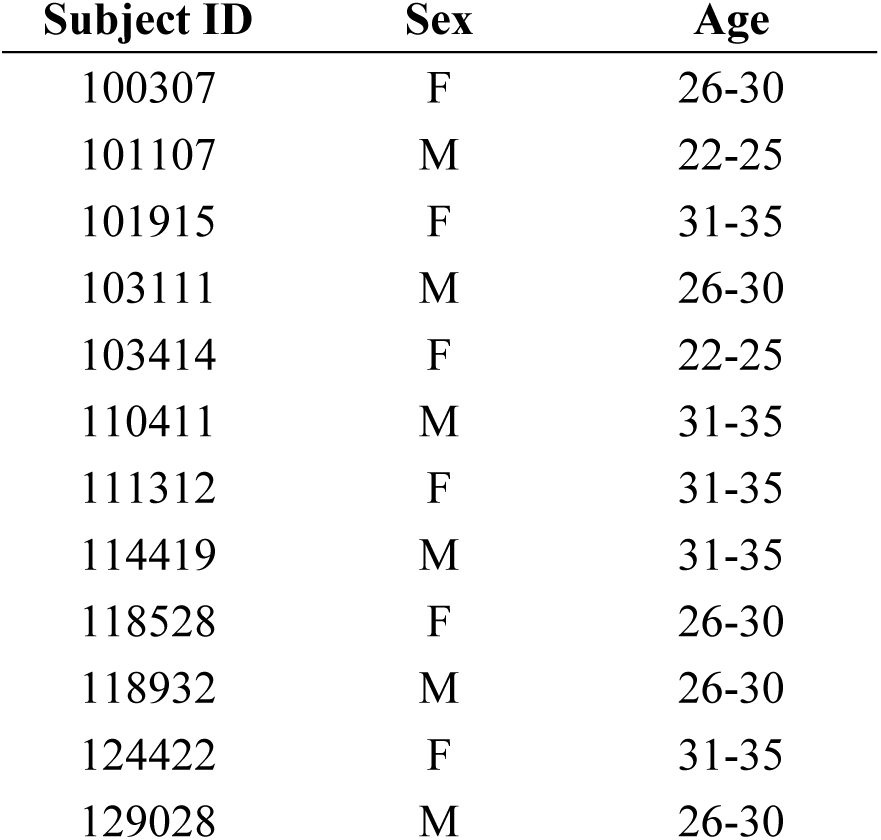
Demographics.

### Neuroimaging protocols and dMRI tractographic analysis

Only 3T data were used for this project. We used minimally processed data, which included voxel-wise correction of diffusion MRI gradients. The T1-weighted (T1W) and dMRI protocols were as follows. The ACPC-aligned T1W MRI images (0.7 × 0.7 × 0.7 mm3 voxel size) from the HCP Young Adult dataset were used for structural MRI analysis. dMRI analysis was performed on the same subjects. The T1-weighted and dMRI protocols were as follows: T1W: 3D MPRAGE, TR = 2400 ms, TE = 2.14 ms, TI = 1000 ms, Flip angle 8°, and voxel size 0.7 mm isotropic. dMRI: Spin-echo EPI, with TR = 5520 ms, TE = 89.5 ms, and flip angle of 78°. The refocusing flip angle was set at 160°. The multifactor was 3, with an echo spacing of 0.78 ms and a voxel size of 1.25 mm isotropic. The b-values used were 1000, 2000, and 3000 s/mm2, each b-shell with 90 diffusion directions and 6 b = 0s. dMRI data preprocessing was conducted using our in-house pipeline (https://github.com/pnlbwh/luigi-pnlpipe). Briefly, diffusion-weighted images were converted to NIFTI, realigned, and re-centered, followed by a semi-automated quality control. Subsequent steps included Gibbs ringing removal (Kellner et al., 2016), brain extraction using a neural-network method (Palanivelu et al., 2020), and correction for motion, eddy currents, and EPI related distortions (Andersson et al., 2003). Diffusion data were analyzed using the Unscented Kalman Filter (UKF) tractography approach (Malcolm et al., 2010; Reddy and Rathi, 2016) as implemented in 3D Slicer (Caviness et al., 1996; Makris et al., 2013). Default settings for UKF were used. Fibers were seeded in voxels with fractional anisotropy (FA) greater than 0.1 (3 seeds per voxel), propagated until FA dropped 0.08 or the mean signal dropped below 0.06. This algorithm uses tractography to drive the local fiber model estimation, that is, model estimation (in this case, the multiple tensors) is done while tracing a “fiber” from seeding to termination.

### Quantitative Analysis

Biophysical parameters of average fractional anisotropy (FA), axial diffusivity (AD) and radial diffusivity (RD) of the fiber tracts in the 12 healthy human subjects for the DA and 5-HT circuits were measured. The presence or absence of specific fiber tracts for all investigated connections of the SN, VTA and DRN in the left and right hemisphere of each individual subject was also recorded. It should be noted that in this project we report on quantitative measures only for the connections that were present in all (12/12 or 100%) or the overwhelming majority (11/12 or 92%) of subjects.

## Results

We were able to delineate the SN and VTA dopaminergic circuits as well as the DRN serotonergic circuits in 12 healthy human subjects of the HCP repository. With respect to the SN and VTA dopaminergic circuits (**Fig. 2A-E**), we delineated the specific connections of these nuclei (**Fig. 2A (SN); 2B (VTA)**) and we also reconstructed the three principal dopaminergic projections, namely the mesostriatal **(Fig. 2C)**, mesolimbic **(Fig. 2D)** and mesocortical **(Fig. 2E)** systems. Regarding the DRN circuitry we delineated its specific connections in the same 12 healthy human subjects of the HCP repository (**Fig. 3**). Although small, this sample provides a pilot dataset of the SN and VTA dopaminergic circuits as well as the DRN serotonergic circuits in humans.

**Figure 2.**
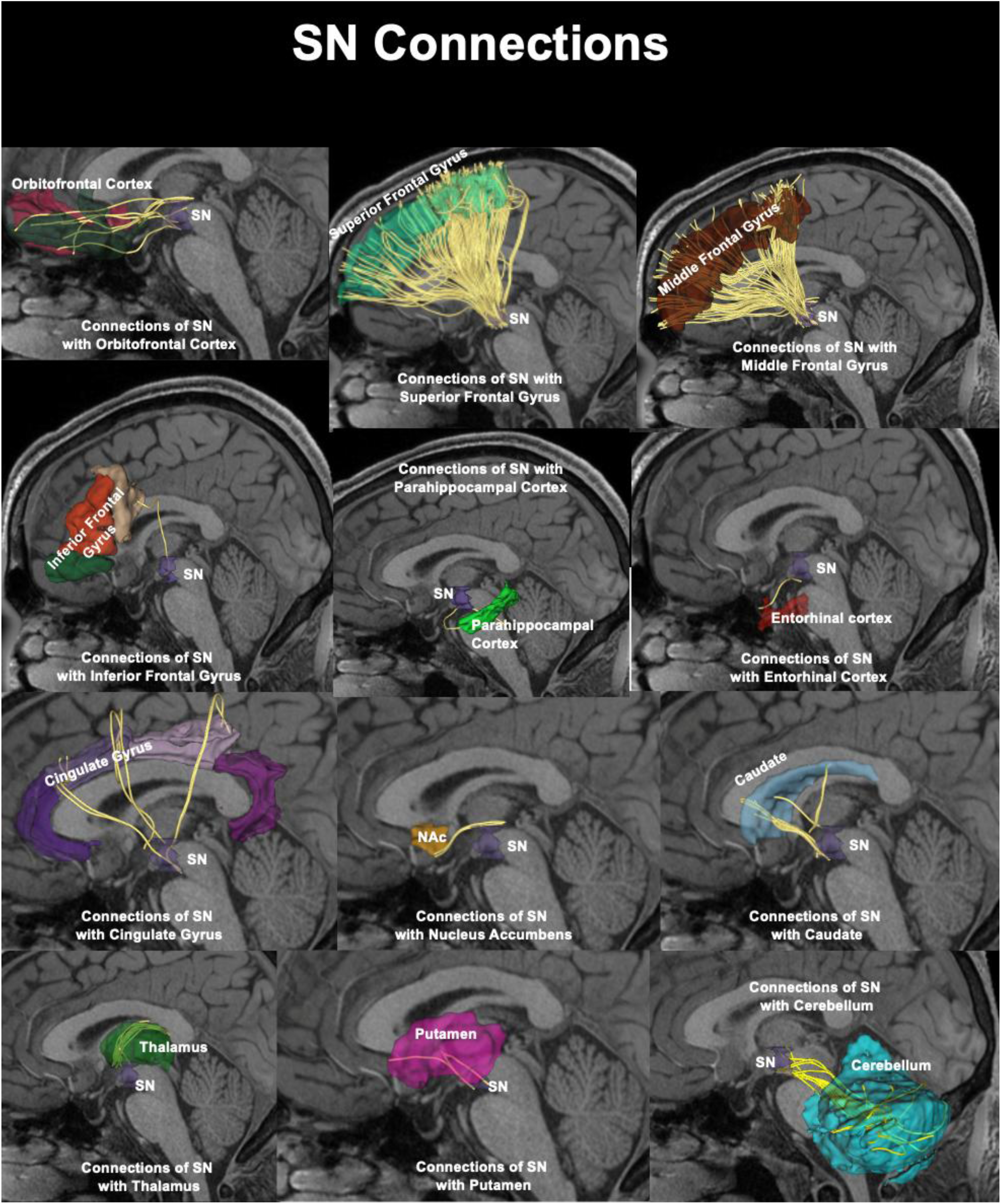
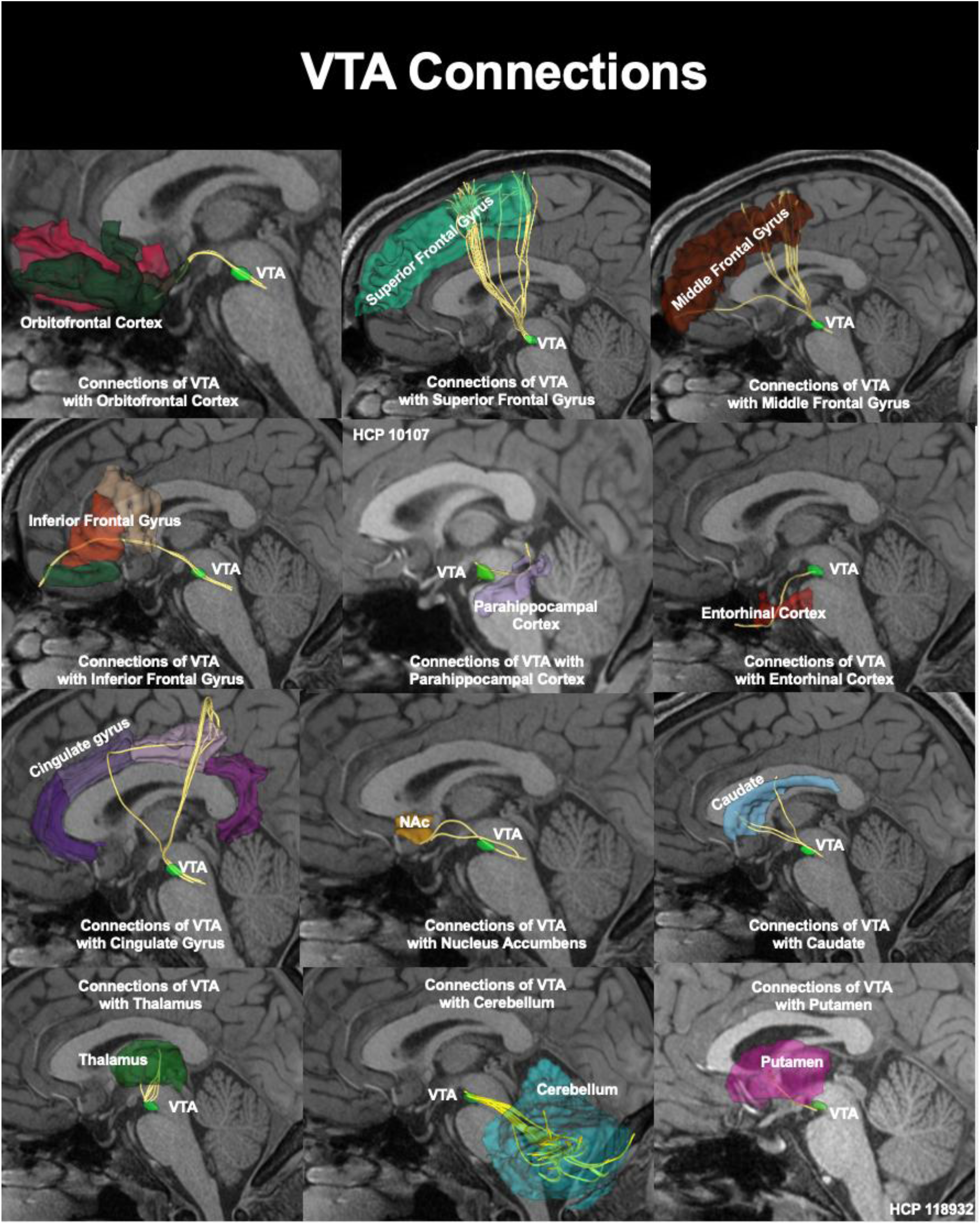
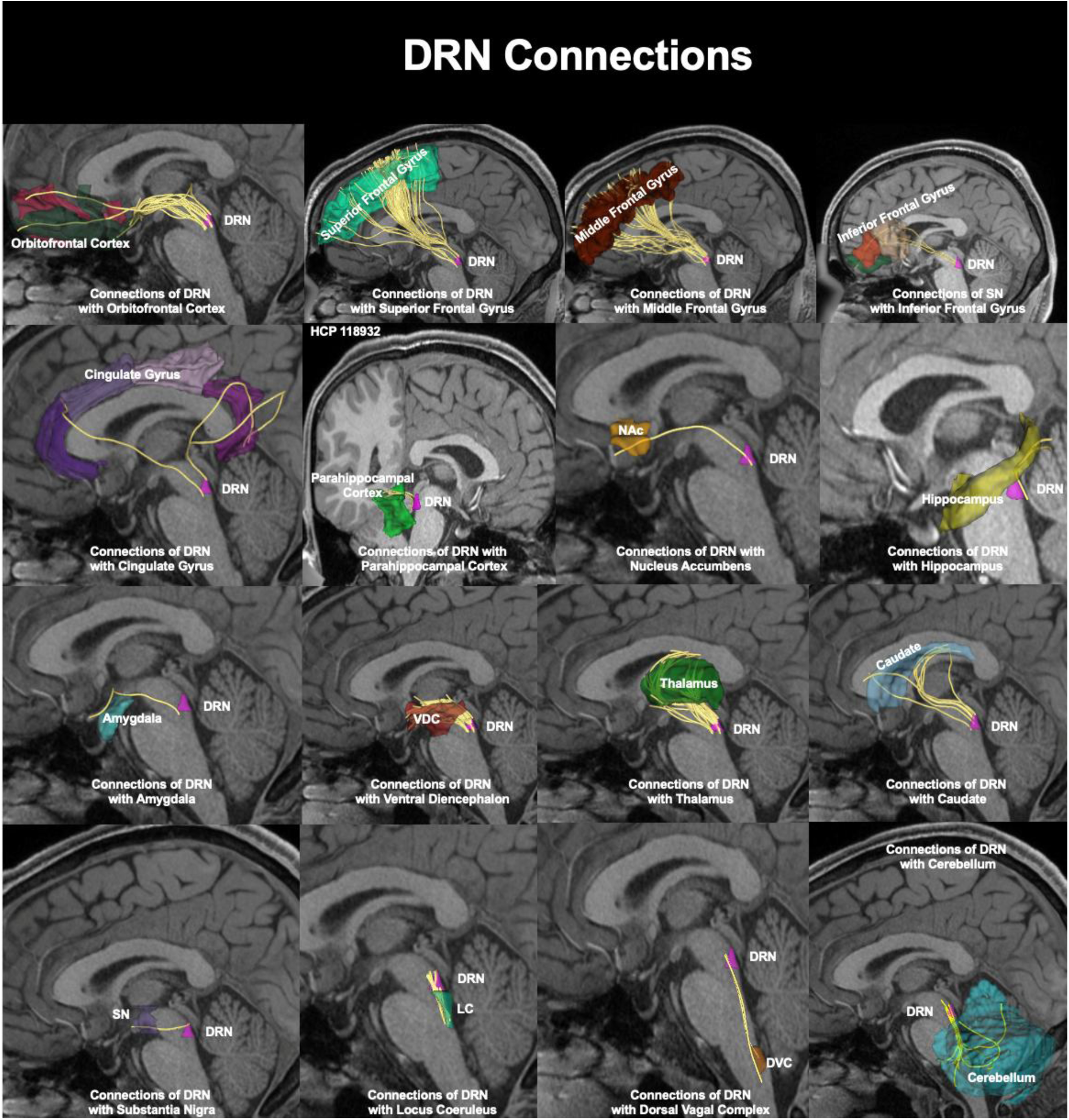
The results of dMRI tractographic analysis are shown for the left side of the brain in a series of representative visualizations. The 12 panels in Figure 2A show the dopaminergic (DA) connections of the substantia nigra (SN) with the orbitofrontal cortex, superior frontal gyrus, middle frontal gyrus, inferior frontal gyrus, parahippocampal gyrus, entorhinal cortex, cingulate gyrus, nucleus accumbens (NAc), caudate nucleus, thalamus, putamen, and cerebellum. Likewise, the 12 panels of Figure 2B show the dopaminergic (DA) connections of the ventral tegmental area (VTA) with the orbitofrontal cortex, superior frontal gyrus, middle frontal gyrus, inferior frontal gyrus, parahippocampal gyrus, entorhinal cortex, cingulate gyrus, nucleus accumbens (NAc), caudate nucleus, thalamus, cerebellum, and putamen. Finally, the 16 panels of Figure 2C show the serotonergic (5-HT) connections of the dorsal raphe nucleus (DRN) with the orbitofrontal cortex, superior frontal gyrus, middle frontal gyrus, inferior frontal gyrus, cingulate gyrus, parahippocampal gyrus, entorhinal cortex and nucleus accumbens (NAc), hippocampus, amygdala, ventral diencephalon (VDC), thalamus, caudate nucleus, substantia nigra (SN), locus coeruleus (LC), dorsal vagal complex (DVC), and cerebellum.

**Figure 3A.**
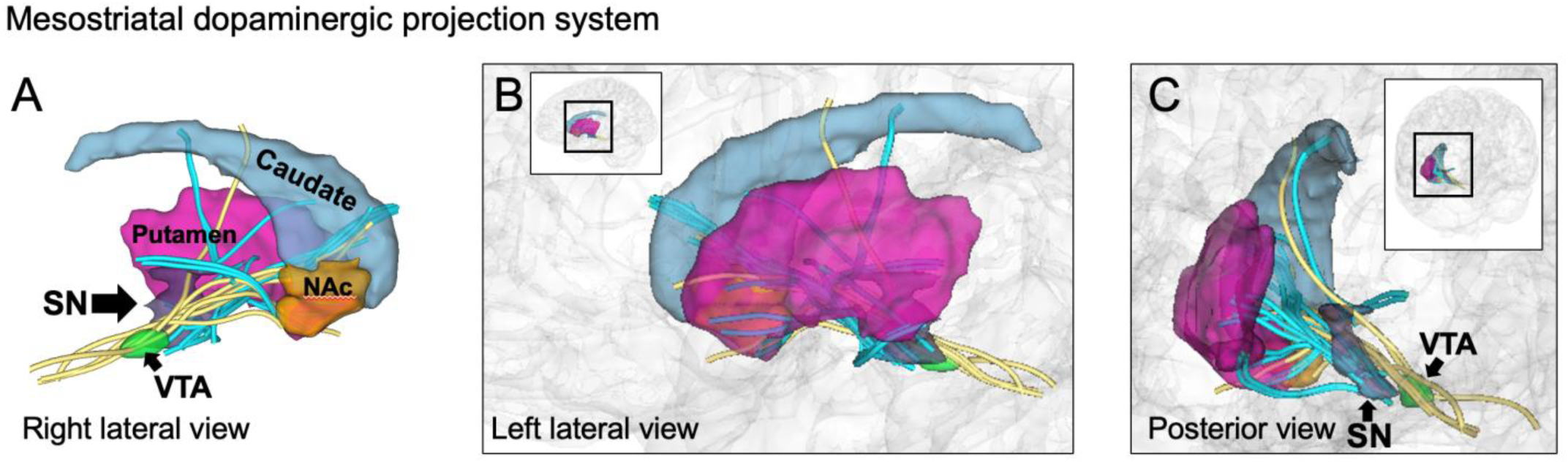
Mesostriatal dopaminergic (DA) projection system involving connections from the substantia nigra (SN) and the ventral tegmental area (VTA) to the caudate nucleus, putamen and nucleus accumbens (NAc), as shown in a right lateral view (A), left lateral view (B) and an oblique posterior view (C). The orientation of B and C are shown at lower magnification in the glass brain insets.

**Figure 3B.**
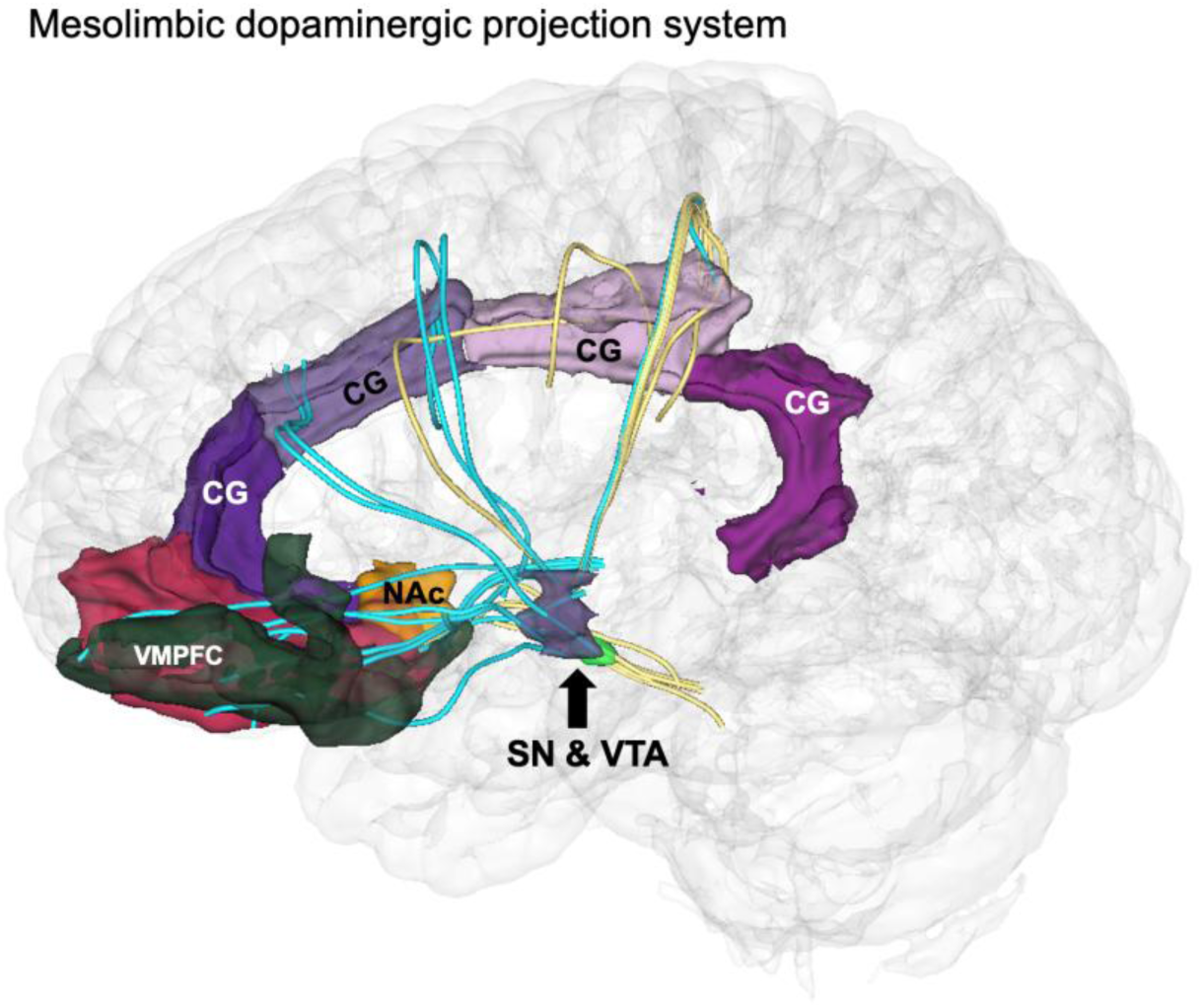
Mesolimbic dopaminergic (DA) projection system involving connections principally from the substantia nigra (SN) and ventral tegmental area (VTA) to the nucleus accumbens (NAc), the components of the cingulate gyrus (CG) and ventromedial prefrontal cortex (VMPFC).

**Figure 3C.**
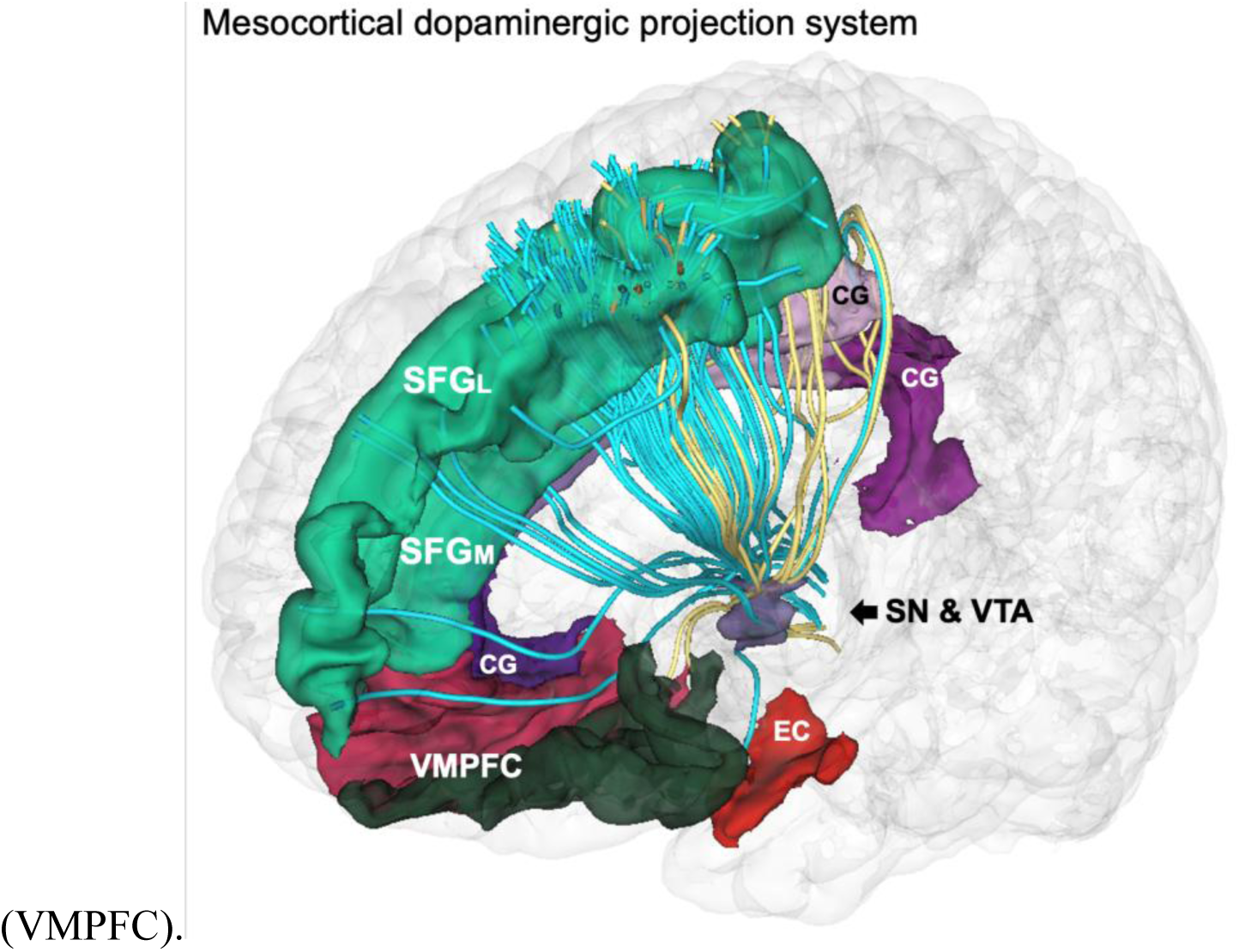
Mesocortical dopaminergic (DA) projection system involving connections from the substantia nigra (SN) and the ventral tegmental area (VTA) to the cingulate gyrus (CG), entorhinal cortex (EC), ventromedial prefrontal cortex (VMPFC) and the medial and lateral portions of the superior frontal gyrus (SFGM and SFGL).Thus, the results of this study provide a) evidence of the SN and VTA dopaminergic circuits and b) evidence of the DRN circuitry in the human brain. Furthermore, the use of a balanced sample of 12 living human subjects from the HCP repository (six females and six males matched for age) provides the basis for a pilot dataset for the human SN, VTA and 5-HT circuits.

Overall, our preliminary results on the DA circuits are proof of principle demonstrating that currently used multimodal clinical neuroimaging allows the detection and delineation of these neurochemical pathways. The sequence of **Figures 2A, 2B, 2C** shows 40 structural connections, namely 12 SN connections **(Fig. 2A)**, 12 VTA connections **(Fig. 2B)** and 16 DRN connections **(Fig. 3C)**. **Figures 3A, 3B** and **3C** show the assembly of individual fiber connections into the three classical projection subsystems of the mesotelencephalic DA system, namely the mesostriatal (**Fig. 3A**), mesolimbic (**Fig. 3B**) and mesocortical (**Fig. 3C**) projections. **Figure 4** shows only the strongest connections that were present with an incidence at or above 92% among the 12 subjects examined in this study. We expect these findings to guide further anatomical investigations in basic and clinical neuroscience.

**Figure 4.**
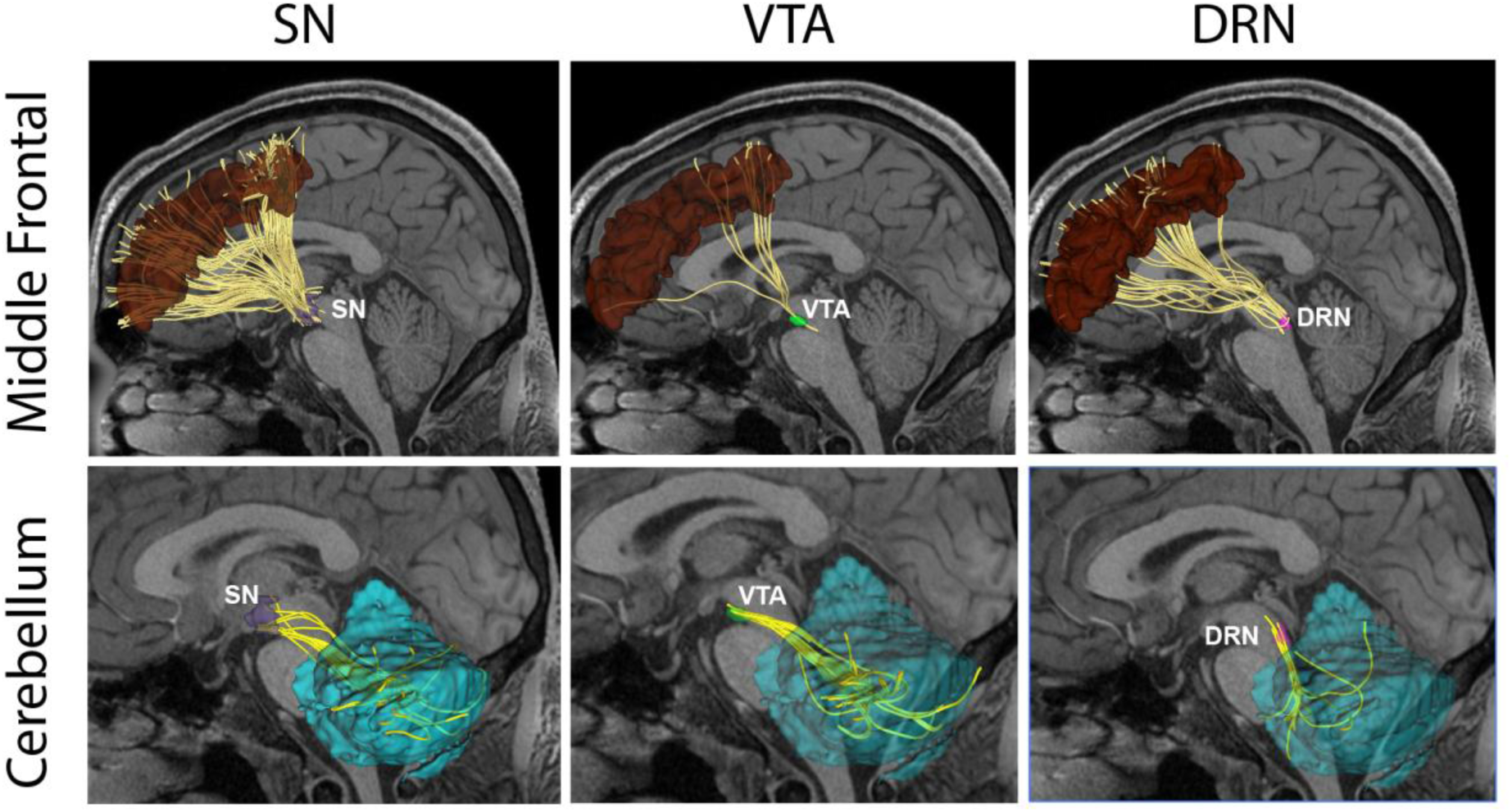
This figure depicts the connections of the SN, VTA and DRN that were present with a frequency of 92% or higher among all subjects. These connections were with the DLPFC (dorsolateral prefrontal cortex, BA46/MFG) and the cerebellum and are highlighted in the shaded columns of **Table 3**.

**Table 3:**
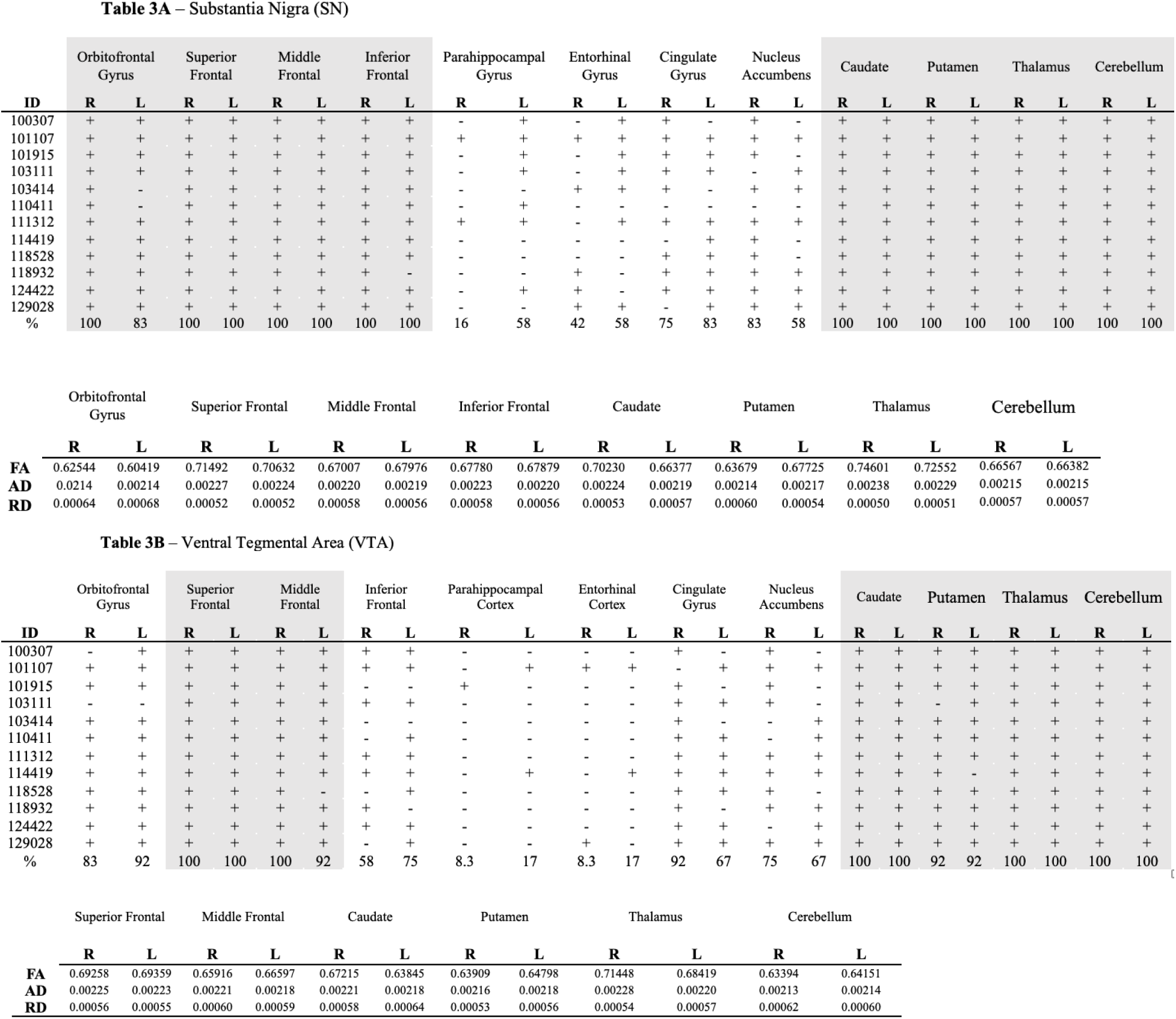
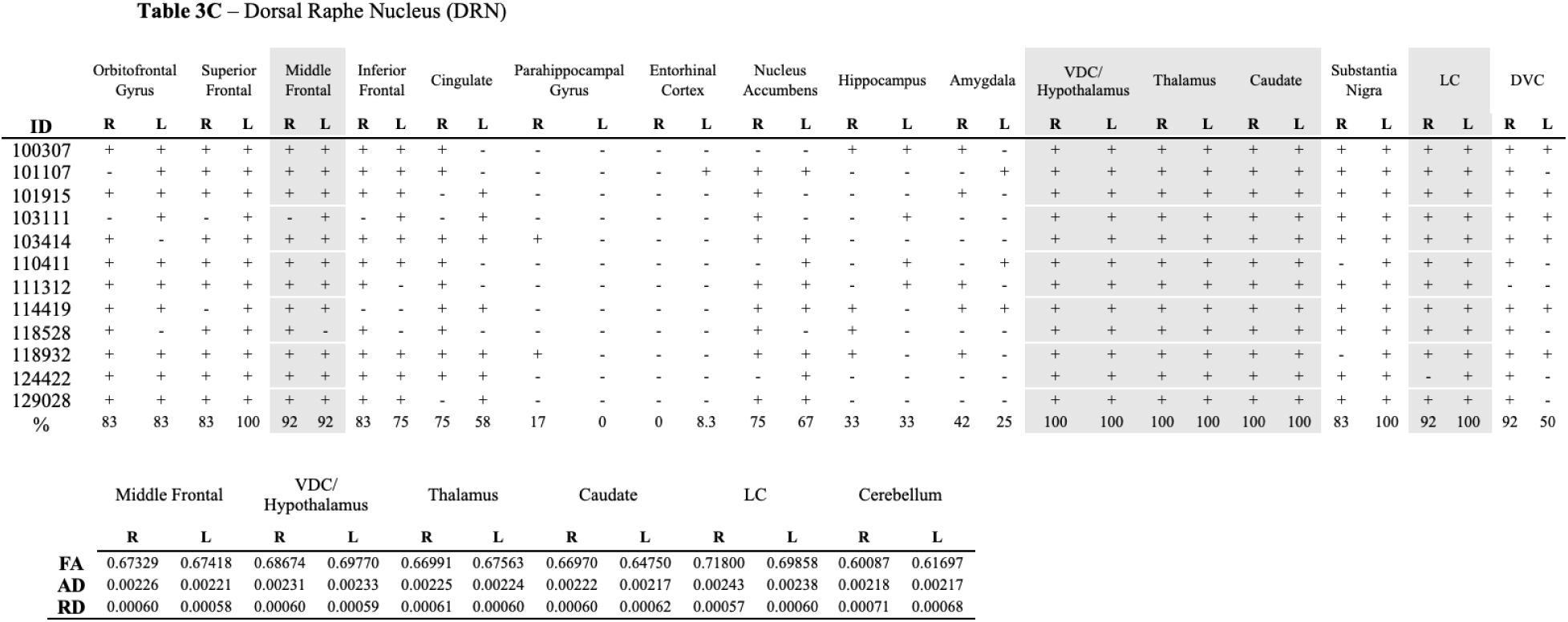
This table shows the presence or absence of specific fiber tracts in the left or right hemisphere of each individual subject. Shaded columns show structural connections present with frequency of 92% or higher. Note that we report the biophysical paramaters of only the fiber tracts that were present with frequency of 92% or more as shown above.

### Quantitative Analysis

The presence or absence of specific fiber tracts in the left or right hemisphere of each individual subject was recorded and the findings are listed in **Table 3** as follows: **Table 3A** shows the results for the SN; **Table 3B** for the VTA; and **Table 3C** for the DRN. The shaded columns indicate connections that were present with a frequency of 92% or higher. Given the importance of noninvasive neuromodulatory treatments currently used in neuropsychiatry, in particular TMS, DLPFC (dorsolateral prefrontal cortex, BA46/MFG) and the cerebellum are highly relevant because of their use as targets. Notably, the DLPFC and cerebellar connections, which are highlighted in **Figure 4**, are among the strongest we observed, specifically the DLPFC at 92% and the cerebellum 100%.

Furthermore, in **Table 3**, the biophysical parameters (i.e., FA, AD and RD) of the structural fiber connections that were present with a frequency of 92% or higher are included separately for the SN (**Table 3A**), VTA (**Table 3B**) and DRN (**Table 3C**). Specifically, the SN fiber connections present with a frequency of 92% or greater were with the orbitofrontal gyrus, superior frontal gyrus, middle frontal gyrus, inferior frontal gyrus, caudate nucleus, putamen, thalamus and cerebellum (**Table 3A**). The VTA fiber connections present with a frequency of 92% or more were with the superior frontal gyrus, middle frontal gyrus, caudate nucleus, putamen, thalamus and cerebellum (**Table 3B**). Finally, the DRN fiber connections present with a frequency of 92% or higher were with the middle frontal gyrus, ventral diencephalon/hypothalamus, thalamus, caudate nucleus, locus coeruleus and cerebellum (**Table 3C**).

### Reliability measurements

Inter-rater reliability estimates were done for two research assistants trained in neuroanatomy and morphometric analysis (PHT, KH) and intra-rater reliability estimates were done by PHT. Inter-rater reliability of the SN, VTA and DRN ROIs was as follows. For the SN the Dice coefficient was SNright = 0.91 and SNleft =0.90 for inter-rater reliability and SNright = 0.90 and SNleft =0.91 for intra-rater reliability. For the VTA the Dice coefficient was VTAright = 0.94 and VTAleft 0.94 for inter-rater reliability, and VTAright = 0.95 and VTAleft = 0.93 for intra-rater reliability. Finally, for the DRN the Dice coefficient was DRNright = 0.99 and DRNleft = 0.99 for inter-rater reliability and DRNright = 0.97 and DRNleft = 0.97 for intra-rater reliability.

## Discussion

In the present study, we report initial results on the human dopaminergic and serotonergic structural circuits. Specifically, we used a novel structural multimodal neuroimaging methodology that combined structural T1-weighted MRI morphometric and dMRI tractographic analyses. In a prior study by our group, we presented evidence on the human coerulean norepinephrine (NE) circuitry (Makris et al., 2024). Importantly, in the present study we generated a pilot dataset for the human brain dopaminergic and serotonergic circuits with balance of age and gender. These data provide quantitative information on the structural circuitry of the three DA systems, i.e., mesostriatal, mesolimbic and mesocortical, which are critically involved in several biobehaviors and clinical conditions such as substance dependence, Parkinson’s disease (PD) and schizophrenia. Likewise, critical connections of the DRN circuitry, such as the DLPFC, caudate, thalamus, and hypothalamus, were clearly delineated. Although preliminary, our findings demonstrate the capability of current clinical multimodal neuroimaging to delineate structural DA and 5-HT circuits in the human brain in normative and clinical conditions.

There has been considerable literature since the 1960s addressing the fiber connections of brainstem nuclei including principally the dopaminergic systems. Whereas the vast majority of these studies have been conducted in animals using traditional experimental methods, there are also several structural connectivity investigations done recently in humans using fiber microdissection (e.g., (Skandalakis et al., 2024) and dMRI approaches (e.g., (Edlow and Wu, 2012; Edlow et al., 2021). The latter studies have been focused mainly on brainstem nuclei in their association with specific behaviors and clinical conditions and are summarized as follows. Edlow and colleagues examined brainstem connections and consciousness, thus considering principally the RAS system and the VTA connections and using a diffusion imaging approach (Edlow and Wu, 2012; Edlow et al., 2021). Several studies have shown connectivity between the SN and dorsal striatum or between the VTA and ventral striatum (i.e., nucleus accumbens septi) using probabilistic dMRI-based tractography (e.g., (Chowdhury et al., 2013; Xie et al., 2013; Handfield-Jones, 2019; Hosp et al., 2019; Rusche et al., 2021). Zhang and colleagues, in particular, showed a tripartite subregional organization of the SN based on its structural connectivity with motor, cognitive and limbic brain regions (Zhang et al., 2017). Cirillo et al. showed connections of SNpc with the thalamus using streamline tractography (Cirillo et al., 2024), whereas other investigations depicted SN and VTA connections with frontal cortical areas (e.g., (Kwon and Jang, 2014; Zhang et al., 2017). Furthermore, Kwon and Jang (2014) reported widespread SN and VTA structural connectivity using probabilistic tractography, and addressed its relevance to diseases such as PD. Menke and colleagues (2010) addressed the question of segmenting the SN into pars compacta (SNpc) and pars reticulata (SNpr) using probabilistic tractography (e.g., (Johansen-Berg et al., 2005)), which is an important issue in assessing the progression of PD (Menke et al., 2010). Trutti and colleagues generated an atlas of the VTA using an MRI-based probabilistic segmentation approach (Trutti et al., 2021). Massey and colleagues studied the SN using histology and showed the great difficulty of separating SNpc from SNpr even with the aid of histology (Massey et al., 2017). Moreover, there have been studies using structural neuroimaging that have shown connectivity between cerebral regions and the brainstem, such as those of Chauvel and colleagues (Chauvel et al., 2023, 2024). Specifically, the comparative study by Chauvel et al. (Chauvel et al., 2023) in the chimpanzee brain has addressed the connectivity between subthalamic and hypothalamic regions and the brainstem. In addition, Coenen and colleagues have reported significant findings in brainstem structural connectivity using neuroimaging, and in nonhuman primates characterized in a more detailed fashion the medial forebrain bundle (MFB). More specifically, Coenen et al. used injection tract tracing methods in the marmoset to determine connections of the ventromedial prefrontal cortex and pre-SMA with diencephalic areas (Coenen et al., 2022). In addition, they corroborated and expanded their studies in human neuroimaging datasets using dMRI tractography (Coenen et al., 2012, 2018, 2022). In these studies, they focused on the MFB showing its relevance in affective behaviors and subdividing it into further components (Coenen et al., 2012). It should be noted that there have been several studies showing the presence of functional connectivity between the cerebellum and VTA using fMRI (e.g., (Nio et al., 2025) as well as between VTA/SN and cerebellum using resting state functional connectivity (rsFCONN) in PD (e.g., (O’Shea et al., 2022)). Notably, Skandalakis et al. (2024) used fiber microdissection to delineate the ventral tegmental area network connecting the VTA with the raphe nuclei, hypothalamus, mammillary bodies, fornix, septal nuclei, nucleus basalis of stria terminalis, nucleus basalis of Meynert, caudate nucleus, putamen, globus pallidus, insula, amygdala, dorsal hippocampus/dentate gyrus, NAc, entorhinal cortex, and prefrontal areas BA10, BA11 and BA12 (Skandalakis et al., 2024). Furthermore, based on their findings they guided the MRI-based tractography in a large number of human imaging datasets and replicated their original microdissection results, thus showing the remarkable potential of this gross anatomy technique to identify and delineate the stems of fiber tracts in the human brain (Skandalakis et al., 2024).

In contrast, there is a striking paucity of structural connectivity studies of the serotonergic circuitry in humans. Bianciardi and colleagues (Bianciardi et al., 2015) provided a template of human brainstem nuclei including the DRN and generated probabilistic labels in the MNI space. Furthermore, Edlow et al. (2016) performed measurements of streamline probability and connectograms of the DRN. More recently, García-Gomar et al. (2022a) showed widespread connections of DRN using automated parcellation approaches, specifically FreeSurfer, for the determination of structural ROIs used as seeds for tractography and probabilistic tractography in combination with graph analysis for the delineation of the DRN structural connectome (García-Gomar et al., 2022a).

It is important to note that studies of the human dopaminergic and serotonergic systems comprise mainly investigations of SN, VTA and DRN connections using probabilistic tractography methods that are limited in their ability to delineate the topographic details and quantitative characteristics of these connections such as FA, AD and RD measures. Moreover, in these prior studies the method of segmentation for the brainstem nuclei, specifically the SN, VTA and DRN, did not explicitly report inter-rater and intra-rater reliability measures, which is a serious methodological confound in anatomical neuroimaging analysis involving segmentation of MRI-based structural ROIs (e.g., (Filipek et al., 1994; Worth et al., 1998; Caviness et al., 1999). It should be noted that there is a large literature on the location and function (including metabolism) of gray matter structures associated with principal neurotransmitter systems of the human brain (e.g., Alzamil, 2025; Shang et al., 2026). Neuroimaging used for this purpose includes such techniques as positron emission tomography (PET) (e.g., Alzamil, 2025; Ceccarini et al., 2020; Hansen et al., 2026; Veldman et al., 2022), single photon emission computerized tomography (SPECT) (e.g., Alzamil 2025; Shang et al. 2026), pharmacological MRI (phMRI) (e.g., Khalili-Mahani et al., 2017; Reneman et al., 2021; Wandschneider et al., 2016) as well as fMRI (including resting-state fMRI, rs-fMRI, and rsFCONN) (e.g., Bruinsma et al., 2018; O’Shea et al., 2022; Saiz-Masvidal et al., 2025). These techniques have contributed significantly to elucidating the anatomical location and distribution of receptors associated with the dopaminergic (e.g., Alzamil 2025; Bruinsma et al., 2018; Ceccarini et al., 2020; Hansen et al., 2026; Khalili-Mahani et al., 2017; O’Shea et al., 2022; Reneman et al., 2021; Saiz-Masvidal et al., 2025; Wandschneider et al., 2016) and serotonergic (e.g., Alzamil 2025; Ceccarini et al., 2020; Hansen et al., 2026; Khalili-Mahani et al., 2017; Saiz-Masvidal et al., 2025; Shang et al. 2026; Veldman et al., 2022; Wandschneider et al., 2016) neurotransmitter systems in the human brain and thus have provided important background information to be used to study clinical conditions. However, given their nature, the aforementioned functional imaging techniques cannot address the specific structure and topography of the anatomical connections between the key structural components of these neurotransmitter systems. This gap in our knowledge can be addressed using dMRI-based tractography, as has been done in the present study.

The present study fills a gap in the literature in the following ways. Firstly, we used a novel structural multimodal neuroimaging methodology that combined T1-weighted MRI morphometric analysis and dMRI WMQL-based tractography. Given its morphometric nature and more precise tractographic analysis, this methodology provides quantitative measures of specific fiber connections between morphometrically specified gray matter regions of interest which are used subsequently as seed ROIs for WMQL-based tractographic analysis. Specifically, these ROIs are determined using a unique morphometric approach, namely brain volumetrics (Caviness et al., 1999), which has inter- and intra-rater reliability measures as a prerequisite condition for the delineation of the ROIs in order to accept the ROIs as valid (e.g., (Filipek et al., 1994; Worth et al., 1998; Caviness et al., 1999). Furthermore, the delineation of fiber tracts was done using dMRI-based tractography, which characterizes structural connections between pre-specified ROIs with clear topographic locations and trajectories. To this end, we used the WMQL, a software tool with improved tractographic capabilities (Wassermann et al., 2013, 2016). We also provided proportions of the presence or absence of these SN, VTA, and DRN fiber connections. Finally, we created a pilot dataset for the most robust, i.e., most reliably present, connections with respect to their biophysical parameters of FA, AD and RD (as shown in **Table 3**). We believe that the fiber pathways detected in the present study at 92% frequency or greater should be reliably detectable in clinical neuroimaging settings. It should be emphasized that structural studies of DA and 5-HT circuits as performed in the present investigation are essential for providing accurate fiber pathway neuroanatomical information in individual brains. Importantly, precise topographic information, which is not provided by probabilistic approaches, can enable clinicians to more accurately and reliably target these brain circuits using invasive and noninvasive neuromodulation interventions, e.g., DBS and TMS, currently used in patient-specific precision neuropsychiatry (see, e.g., (Makris et al., 2016). Furthermore, the monitoring and assessment of neuropsychiatric interventions could become more precise, complete and informative once the structural neuroanatomy of these neurotransmitter systems is determined and integrated with functional information from other methods such as functional neuroimaging and behavioral-clinical assessments. Moreover, precise delineation of specific fiber tracts and measurement of their biophysical parameters allows the detection and assessment of pathological processes such as neuroinflammation or demyelination affecting those fiber tracts (e.g., (Pasternak et al., 2016; Kikinis et al., 2024). Such detailed anatomical information could enable clinicians using MRI to monitor a pathological process as well as the efficacy of a therapeutic intervention over time.

The present study differs from other neuroimaging dMRI tractographic studies conceptually and methodologically as well as in terms of results. For example, the study by Chauvel and colleagues (Chauvel et al., 2024) as well as those by Coenen and colleagues (Coenen et al., 2012, 2018, 2022) followed a traditional neuroanatomical rationale and methodological approach that did not focus on specific human neurochemical pathways as in the present study. Instead, these authors followed a coarse neuroanatomical approach involving multiple structures and connectional pathways rather than delineating individual connections of brainstem nuclei associated with specific neurochemical systems. Thus, our approach differs from these studies in a number of important ways. Fundamentally, we followed a quantitative approach of brain morphometry, which takes into serious consideration the neuroanatomical individuality, and thus the biological structural variability, of an individual’s brain, generating methods that are valid in terms of their anatomical accuracy and reliability (e.g., (Caviness et al., 1996; Makris et al., 2005). Thus, the definition of brain regions of interest (ROIs) is based on landmarks that are reliably identifiable by a certain neuroimaging modality. Following this approach, all ROIs, such as the SN, VTA, and DRN, that represent the origins and terminations for the determination of specific dopaminergic and serotonergic fiber pathways were parcellated. We then used WMQL, which enhances the accuracy of connectional data by restricting tractography between the pre-specified origin and termination ROIs. Thus, the current study clearly advances our critical knowledge of the structural neuroanatomy of the human dopaminergic and serotonergic brain circuits. Importantly, it indicates the potential of current structural multimodal neuroimaging to study systematically the neurochemical systems in the human brain, a topic of great relevance in basic and clinical neuroscience. Our approach to delineating specific neurochemical systems of the human brain will contribute to the finer elucidation of complex, multi-component fiber pathways such as the MFB. As has been pointed out by Swanson, the MFB is considered “the most complex of the longitudinal central nervous system white matter tracts” (Swanson, 2015, p. 430). A thorough and extensive neuroanatomical literature review of the MFB and other neurochemical pathways of the brain is beyond the scope of the current discussion. Given the relevance of neurochemical circuits in current neuropsychiatry, we outline below the neuroanatomy of the dopaminergic and serotonergic circuits.

### DA circuitry

Dopamine (DA) neurons of origin and DA production are located mainly in the brainstem and hypothalamus. Cell hubs producing DA in the cerebrum (or telencephalon) are found in the olfactory system and the outer zone of the olfactory bulb in particular, which is referred to as the A16 cell group (Halász et al., 1977; Priestley et al., 1979). The connections of the DA system follow a medial (or “meso”) trajectory within the neuraxis and extend within and between different subdivisions of the CNS. Given its topography and structural connectivity, the DA system was originally labeled the “mesotelencephalic dopaminergic system” referring mainly to the entire DA forebrain projection of its midbrain cell groups of origin (Fallon and Moore, 1978; Moore and Bloom, 1978; Lindvall et al., 1983). This massive ascending mesencephalic projection, coursing principally from cell groups A8 (LTA), A9 (SNpc) and A10 (VTA), consists of two principal systems that were named by Ungerstedt (Ungerstedt, 1971) as the “nigrostriatal system” and the “mesolimbic system”. Eventually, Nieuwenhuys (1985) proposed a more practical typology by subdividing the DA system into three principal projections and seven additional ancillary connections as follows: a) the “mesostriatal projection”, coursing from the LTA (A8 cell group), SNpc (A9 cell group) and VTA (A10 cell group) (e.g., (Anden et al., 1964; Moore and Bloom, 1978; Lindvall et al., 1983); b) the “mesolimbic projection”, originating from the VTA (A10 cell group) (3); and c) the “mesocortical projection”, arising from the VTA (A10 cell group) and the medial portion of the substantia nigra. The mesostriatal projection connects the SNpc (A9 cell group) principally with the caudate nucleus and putamen. It also connects the VTA (A10 cell group) mainly with the nucleus accumbens septi (NAc), and the LTA (A8 cell group) with the ventral putamen (Hökfelt et al., 1974; Emson and Koob, 1978; Lindvall and Björklund, 1978; Nieuwenhuys, 1985). The mesolimbic projection connects the VTA with the NAc and ascends within the medial forebrain bundle (MFB), in a location medial to the mesostriatal projection. It also connects the VTA with olfactory areas such as the olfactory bulb and tubercle and the anterior olfactory nucleus. Furthermore, it connects the VTA with the anterior perforated substance, the lateral septal nucleus, the bed nucleus of the stria terminalis and, importantly, with the central and basal nuclei of the amygdala. The mesocortical projection originates from the VTA and the medial portion of the substantia nigra, connecting with the medial aspect of the frontal lobe in each cerebral hemisphere, namely the piriform and pre-piriform olfactory cortices, the entorhinal cortex and, notably, the anterior cingulate cortex (ACC) (Hökfelt et al., 1974; Emson and Koob, 1978; Lindvall and Björklund, 1978; Nieuwenhuys, 1985).

### Serotonergic circuitry

Serotonin (5-HT) neurons of origin are situated mainly in the brainstem. 5-HT is produced principally by the B cell groups of neurons in the raphe nuclei (e.g., (Taber et al., 1960; Braak, 1970; Nieuwenhuys, 1985) located in the median and paramedian zones of the medulla, pons and midbrain. It should be noted that the B cell groups are co-localized to a considerable extent with the raphe nuclei. The raphe nuclei are multiple neurotransmitter complexes, which include neurons associated with the synthesis of neurotransmitters in addition to 5-HT (Wiklund and Björklund, 1980; Bowker et al., 1983). There are nine distinguishable cell groups, namely B1 to B9 (Dahlström and Fuxe, 1964). Specifically, the B1 and B2 cell groups are located in the medulla and coincide, respectively, with the nucleus raphe pallidus and nucleus raphe obscurus. The B3 cell group coincides to a considerable extent with the nucleus raphe magnus at the level of the lower pons. Cell group B4 is located in the upper medulla in proximity with the medial vestibular nuclei, whereas cell group B5 is co-localized with the nucleus raphe pontis at the mid-pontine region. The B6 and B8 cell groups are situated mainly, but not exclusively, within in the upper pons in the nucleus centralis superior (of Bechterew) (Sladek and Walker, 1977; Takeuchi et al., 1982). B9 is a small cell group located ventral to the superior central nucleus in the vicinity of the medial lemniscus (Felten and Sladek, 1983). Finally, and importantly for the present study, the B7 cell group is a large neuronal group localized principally at the level of the uppermost pons and lower midbrain within the dorsal raphe nucleus (DRN). An important structural characteristic of the serotonergic neurons is that they are highly arborized, with rich branching of their fibers. This morphological feature accounts for their massive innervation of the brain and makes them the most expansive neurotransmitter system in the central nervous system (CNS). In fact, most of the serotonergic innervation of the brain including the cerebral cortex, thalamus, hypothalamus and basal forebrain as well as the amygdala and the entire brainstem arises from the DRN (Monti, 2010). **Figure 5** illustrates the widespread nature of the DRN serotonergic innervation in the brain of one of the HCP subjects analyzed in the present study (**HCP ID#124422**).

**Figure 5.**
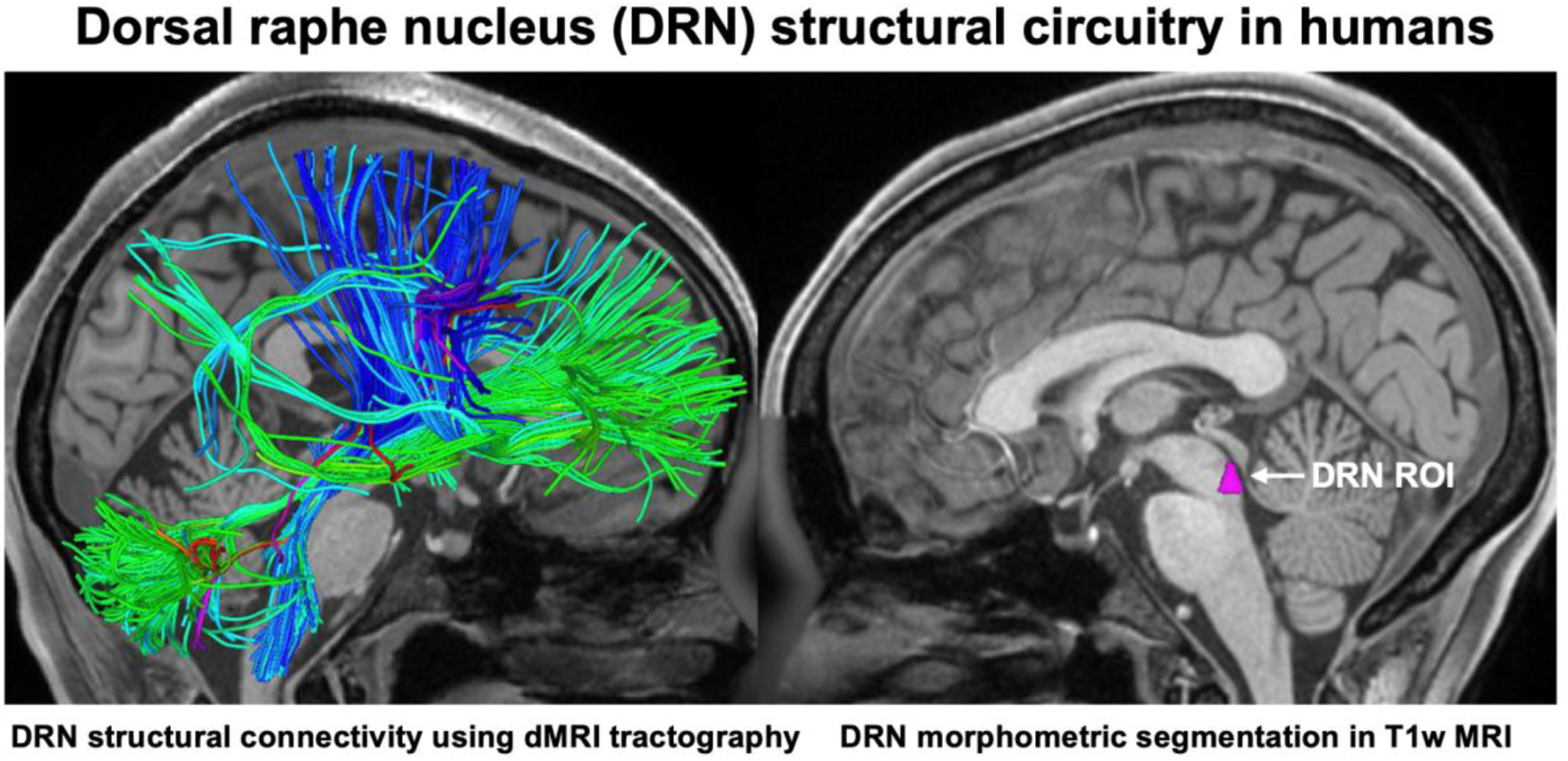
Widespread connections of the DRN serotonergic circuitry with the cerebrum, brainstem and cerebellum in the brain of one of the HCP subjects analyzed in the present study (**HCP ID#124422**).

The axons of serotonergic neuronal groups principally form three distinct fiber pathways as follows: a) The ventral ascending serotonergic pathway, which is one of the largest contributing fiber tracts to the medial forebrain bundle (MFB), connects the B7 cell group and other serotonergic groups such as B6 and B8, with the hypothalamus (e.g., (Moore, 1977). b) The dorsal ascending serotonergic pathway also stems from the B7, B6 and B8 cell groups as well as from the B3 cell group (e.g., (Nieuwenhuys, 1985). Its axonal fibers contribute to the dorsal longitudinal fascicle of Schütz (e.g., (Bobillier et al., 1976; Moore et al., 1978; Parent, 1981) and connect these serotonergic cell groups with the hypothalamus and mesencephalic central or periaqueductal gray (PAG). c) The cerebellar serotonergic structural connections arise from all raphe nuclei including DRN (e.g., (Pierce et al., 1977) and their connections are with both the deep nuclei and the cortex of the cerebellum (e.g., (Takeuchi et al., 1982). Finally, there are several serotonergic connections with other brainstem nuclei, such as the SN, locus coeruleus (LC), parabrachial nucleus (PBN), dorsal motor nucleus of the vagus (DMN) and reticular formation (RF) (e.g., (Nieuwenhuys, 1985).

From this brief neuroanatomical outline, it appears that the MFB and its division into broad subcomponents is an oversimplification of the brainstem structural connectivity and by no means reflective of the specific anatomical architecture of the brainstem neurochemical systems. As discussed above, in a recent paper Chauvel and colleagues (Chauvel et al., 2024) delineated the brainstem white matter connections as a whole, incorporating within one large white matter pathway-like structure dozens of individual fiber tracts by labeling it the diencephalic-mesencephalic pathway (Chauvel et al., 2024). By contrast, Coenen and colleagues (Coenen et al., 2012, 2018, 2022) followed a traditional anatomically driven approach and incorporated into the MFB almost the entire brainstem connectivity with the diencephalon and telencephalon (Coenen et al., 2012, 2018, 2022). Thus, the observations by Chauvel and colleagues and Coenen and colleagues are fundamentally different from those produced by the approach followed in the present study. Specifically, we worked from both a neuroanatomical and a neurochemical perspective at the same time, establishing the structural connectivity of the dopaminergic and serotonergic neurochemical systems. To this end, we adopted a manual morphometric approach to segment, as accurately as possible, specific brainstem nuclei, namely the SN, VTA and DRN, and cerebral ROIs and used a novel dMRI tractography technique, namely the WMQL, which enables the specification of origin and termination of virtual connections between ROIs. Conceptually, our approach is neurochemically driven and more granular, meaning that we delineated circuits of specific neurochemical ROIs such as the SN, VTA and DRN, instead of sampling holistically fiber tracts that contain multiple structural pathways and neurochemical systems. Specifically, multi-component fiber tracts such as MFB, which have recently been sampled in a unitary fashion because of the spatial resolution limitations of MRI morphometric and dMRI tractographic analysis, need to be delineated individually bearing in mind that their various components have distinct origins and terminations and potentially distinct functions. Following this more granular approach, we were able to relate our findings on dopaminergic and serotonergic pathways in the human brain with established neurochemical pathways in nonhuman brains.

### Functional and clinical roles of DA and 5-HT systems

Depending on the specific circuit, the dopaminergic (DA) system manifests different functional roles. The mesostriatal DA projection provides the ability to switch motor programs (Cools, 1980; Cools et al., 1984; Jaspers et al., 1984), which is a highly valuable behavior for a biological organism to operate effectively under constantly changing environmental scenarios. This ability to flexibly switch motor programs extends to cognitive functions as well, which is especially relevant in humans. Dopamine, in conjunection with other neurotransmitters, serves motor behavior. In particular, dopamine appears to coordinate the voluntary, or pyramidal, and involuntary, or extrapyramidal, systems. Dopamine deficit is considered the principal cause of Parkinson’s disease, a movement disorder characterized by uncontrolled movements, stiffness, slowing of movement and difficulty with balance that raises the risk of falls. Changes in dopamine levels have been associated with other clinical conditions such as restless legs syndrome and attention deficit hyperactivity disorder (ADHD). The mesolimbic DA projection of the dopaminergic system plays a role in the experience of pleasure and reward as well as in many other functions, including, e.g., memory, motivation, mood, attention and movement. This is reflected in the connections of the SNpc with the NAc, which are strongly related to reward functions and are considered as the principal “reward center” in the brain (e.g., (Breiter et al., 1997; Berridge, 2007; Koob and Volkow, 2010). The reward network may be an integral part of the neurobiology of drug addiction (Koob, 1999). Structural alterations in this system may enhance the risk for reward deficiency syndrome (Blum et al., 1996; Bowirrat and Oscar-Berman, 2005) and drug-seeking behaviors. This circuitry seems to be critical to abnormalities in hedonic set points associated with increased drug use and drug dependence (Koob, 1992, 1999). By contrast, when there is a decrease in dopamine, depression-like symptoms may appear, such as apathy or feelings of hopelessness. Furthermore, hyperactivity of the VTA and its dopaminergic outflow to limbic structures may play a role in schizophrenia (Stevens, 1973; Bird et al., 1979; Stevens et al., 1979; Reynolds, 1983).

The serotonergic system is related to a range of functions depending on the specific brainstem cell groups and their associated circuits. In general, the DRN circuitry is involved with behaviors such as mood regulation, sleep and sexual activity. Importantly, this circuitry has a key role in cognitive processes such as attention, memory and executive functions including inhibitory control (e.g., (Wingen et al., 2008; Enge et al., 2011; Cowen and Sherwood, 2013; Pattij and Schoffelmeer, 2015; Aznar and Hervig, 2016; Weinberg-Wolf et al., 2018; Coray and Quednow, 2022; Colwell et al., 2024). This array of functions is subserved by the rich structural connectivity of the DRN as demonstrated by the results of this study. Indeed, the widespread circuits linking DRN with the cerebral cortex, in particular the DLPFC and lateral parietal cortical regions, could explain the serotonergic influence on executive functions and attention. By contrast, DRN innervation of the cingulate gyrus and other limbic and autonomic structures could be associated with mood regulation, sleep (Edlow and Wu, 2012; Edlow et al., 2012, 2021) and sexual activity. pathophysiology of these circuits and their dysfunction is reflected in several pathologies. 5-HT is considered to affect development and has been implicated in neurodevelopmental disorders such as autism (e.g., (Sodhi and Sanders-Bush, 2004; Muller et al., 2016; Andersson et al., 2021; Wegiel et al., 2024). Among the known health conditions affected by serotonergic deficits are mood disorders, such as major depressive disorder (e.g., (Bremshey et al., 2024; Shu et al., 2025), suicidal behavior (e.g., (Mann et al., 1990; Underwood et al., 2018; Lagerberg et al., 2022), anxiety disorder (e.g., (Heesbeen et al., 2024; Tseilikman et al., 2024); sleep problems ((Monti, 2011; Wilson et al., 2018; Aung et al., 2024), OCD (e.g., (Charney et al., 1988; Derksen et al., 2020; Kim et al., 2020), PTSD (e.g., (Zhao et al., 2017; Seidemann et al., 2021; Ogłodek, 2022) and schizophrenia (e.g., (Quednow et al., 2020; Schulz et al., 2022; Canul-Medina et al., 2024; Jiménez-Trejo et al., 2024). Recently, the possible involvement of 5-HT brain circuitry in brain-immune interactions and its relationship with acute COVID-19 and long COVID fatigue as well as long COVID cognitive impairment further highlight the intricate connection between the immune system and neurotransmitter regulation (e.g., (Eslami and Joshaghani, 2024; Kikinis et al., 2024).

Overall, by integrating information on dopaminergic and serotonergic circuitry architecture with traditional morphometric analysis in the human brain using current T1-weighted MRI and dMRI we have pioneered an approach which we expect to be of significant value in both basic and clinical neuroscience. It should be noted that most medication-based treatments in neuropsychiatry make use of neurotransmitters. Specific anatomical knowledge of dopaminergic and serotonergic circuitry is critical for targeting these systems and assessing their response to treatment using neuroimaging in neuropsychiatric conditions as elaborated above. Importantly, noninvasive neuromodulation therapeutics such as TMS will benefit from a more precise knowledge of potential target structures such as the SN, VTA and DRN via their direct connections with other brain regions, particularly the cerebellum (see **Fig. 4** and **Table 3**), in ad hoc therapeutic interventions.

### Limitations and future studies

Brainstem neuroanatomical investigations using neuroimaging are currently a topic of great interest but present several limitations. T1- and T2-weighted MRI-based morphometric approaches for in vivo investigations in human subjects, as used in the present study, lack the necessary spatial resolution and signal-to-noise ratio to make them comparable to neuroanatomical and histopathological observations. This is especially the case in brain regions such as the SN, VTA and DRN as well as several other structures in the brainstem, which are characterized by heterogeneous and complex anatomical architecture. Likewise, dMRI tractography studies can be only approximate compared to the tract tracing observations derived from experimental animal studies. More specifically, because heteromodal association areas are greatly expanded in humans as compared to nonhuman primates, homology between monkey and human is stronger for non-cerebral cortical structures such as the brainstem; in general any connections beyond the brainstem and hypothalamus are expected to have less overlap. Our understanding in this regard will become clearer by using T1/T2-weighted MRI and ultra-high resolution dMRI datasets, which have not been used in the present study. The use of only a single MRI modality can lead to serious errors in anatomical interpretation. Investigating structural connections in particular bears the risk of erroneous observations of origins and terminations of fiber tracts that are not well known from the human literature; thus, the ROIs of origin and termination of fiber tracts need to be reliably segmented prior to tractography. In sum, there is a necessity to use more than one MRI modality to minimize inaccuracy when using current methods. It should be noted that much of our knowledge of structural connectivity of the human brain has been derived principally from other animals, such as rodents, cats, dogs or nonhuman primates, as has been elaborated upon in previous reports by our group and others (e.g., (Rushmore et al., 2020)). Taking into account these key considerations, we investigated structures that we were able to identify reliably in our MRI-based studies of the human brain (e.g. (Caviness et al., 1996; Makris et al., 2005)). Furthermore, because the contralateral structural connectivity of the SN, VTA and DRN remains largely unexplored in experimental animals and humans, we focused the present study on ipsilateral structural connectomes of these three neurochemical systems (e.g., (Nieuwenhuys, 1985)). Herein we demonstrated that it is possible to delineate several fiber tracts associated with the dopaminergic and serotonergic systems in humans using current neuroimaging techniques. Notably, we were able to demonstrate reliably several connections of the SN, VTA and DRN with brain regions such as the DLPFC and cerebellum with a frequency of 92% or greater among all subjects. With respect to several other connections the results were less consistent. We believe that the lower rates of presence for certain fiber pathways may be related in part to the complex architecture of the white matter between the brainstem, the diencephalon and the cerebral cingulate, caudate nucleus and putamen regions. This difficulty could be attributed to the massive number of white matter fibers, e.g., of the anterior limb of the internal capsule and the corticospinal tract, that connect the cerebrum with the brainstem and spinal cord. In conclusion, this study demonstrates the critical importance of both in-depth neuroanatomical knowledge and expertise in multimodal neuroimaging analysis. Our current findings for the SN and VTA dopaminergic system as well as the DRN serotonergic system are highly encouraging for studying these structures clinically using MRI. Nevertheless, further technological advances are needed to produce neuroimaging protocols with optimal voxel size and increased signal-to-noise ratios to enable a more refined understanding of these pathways in the human brain. It should be noted that in this project we reported on quantitative measures only for the connections that were present in all (12/12 or 100%) or the overwhelming majority (11/12 or 92%) of subjects. Since this dataset is small and thus represents preliminary data, any follow-up structural connectivity study should include a larger number of subjects and explore the sensitivity and specificity of our method. Furthermore, given that our sample size is limited and that this is a pilot study, we report only descriptive measures recognizing that future studies with larger samples should address issues such as individual variability of these circuits as well as sex differences, developmental differences and hemispheric lateralization. Finally, future studies should include multimodal neuroimaging combining structural as well as functional (including metabolic) techniques to more effectively assess neurotransmitter systems in the human brain, a matter of great relevance in clinical settings.

## 5. Conclusions

We used a novel structural multimodal neuroimaging methodology that combined structural T1-weighted MRI morphometric and dMRI tractographic analyses to delineate the human dopaminergic and serotonergic circuits. The present results provide preliminary evidence for the capability of current multimodal clinical neuroimaging to map the dopaminergic and serotonergic circuits related to SN, VTA and DRN. Among the strongest connections of the SN, VTA, and DRN were those with the DLPFC and the cerebellum, a remarkable finding given the relevance of the DLPFC and the cerebellum, e.g., as putative TMS targets in clinical neuromodulation approaches. The present pilot dataset also demonstrates the potential for generating a quantitative normative database of these circuits using current neuroimaging techniques. Quantitative structural imaging studies of the dopaminergic circuitry of the SN and VTA can be used to advance our understanding of several disease states, such as Parkinson’s disease, ADHD, schizophrenia and drug addiction. Similarly, quantitative structural neuroimaging studies of the DRN serotonergic circuitry may lead to a greater understanding of clinical conditions such as mood disorders, OCD, anxiety disorders, PTSD, sleep problems, COVID-19, as well as autism and schizophrenia. We expect the present neuroanatomical research to encourage further technological development in neuroimaging that will empower clinical applications with the use of MRI protocols with higher signal-to-noise ratios and higher spatial resolution. Furthermore, the monitoring and assessment of neuropsychiatric interventions could become more complete and informative once the structural neuroanatomy of these systems is determined and integrated with information from other methods such as functional neuroimaging and behavioral-clinical assessments.

## 6. Patents

Not Applicable.

### Author Contributions

All authors have read and agreed to the published version of the manuscript. Conceptualization: Nikos Makris and Agustin Castañeyra-Perdomo; Investigation: Nikos Makris, Poliana Hartung Toppa, Richard J. Rushmore, Kayley Haggerty, George Papadimitriou, Agustin Castañeyra-Perdomo; Methodology: All authors; Writing – original draft: All authors; Writing – review and editing: All authors.

### Funding

We would like to acknowledge the following grants for their support: R01MH112748 (NM, RJR), R01AG042512 (NM), K24MH116366, R01MH132610, R01MH125860 (NM), and R21NS136960 (RJR, NM), R01NS125307 (RJR, NM).

### Institutional Review Board Statement

Not Applicable.

### Informed Consent Statement

Not applicable.

### Data Availability Statement

Data were provided in part by the HCP, WU-Minn Consortium (Principal Investigators: David Van Essen and Kamil Ugurbil; 1U54MH091657) funded by the 16 NIH Institutes and Centers that support the NIH Blueprint for Neuro-science Research, and by the McDonnell Center for Systems Neuroscience at Washington University. https://www.humanconnectome.org/study/hcp-young-adult/document/hcp-citations

### Conflicts of Interest

The authors declare no conflicts of interest. The funders had no role in the design of the study; in the collection, analyses, or interpretation of data; in the writing of the manuscript; or in the decision to publish the results.

## Notes

### Competing Interest Statement

The authors have declared no competing interest.

